# Location and concentration of aromatic-rich segments dictates the percolating inter-molecular network and viscoelastic properties of ageing condensates

**DOI:** 10.1101/2022.12.14.520383

**Authors:** Samuel Blazquez, Ignacio Sanchez-Burgos, Jorge Ramirez, Tim Higginbotham, Maria M. Conde, Rosana Collepardo-Guevara, Andres R. Tejedor, Jorge R. Espinosa

## Abstract

Maturation of functional liquid-like biomolecular condensates into solid-like aggregates has been linked to the onset of several neurodegenerative disorders. Low-complexity aromatic-rich kinked segments (LARKS) contained in numerous RNA-binding proteins can promote aggregation by forming inter-protein *β*-sheet fibrils that accumulate over time and ultimately drive the liquid-to-solid transition of the condensates. Here, we combine atomistic molecular dynamics simulations with sequence-dependent coarse-grained models of various resolutions to investigate the role of LARKS abundance and position within the amino acid sequence in the maturation of condensates. Remarkably, proteins with tail-located LARKS display much higher viscosity over time than those in which the LARKS are placed towards the center. Yet, at very long timescales, proteins with a single LARKS—independently of its location—can still relax and behave as high viscous liquids. However, phase-separated condensates of proteins containing two or more LARKS become kinetically trapped due to the formation of percolated *β*-sheet networks that display gel-like behaviour. Furthermore, as a work case example, we demonstrate how shifting the location of the LARKS-containing low-complexity domain of FUS protein towards its center effectively precludes the accumulation of *β*-sheet fibrils in FUS-RNA condensates, maintaining functional liquid-like behaviour without ageing.

## I. INTRODUCTION

Eukaryotic cells achieve spatiotemporal organization of their many biomolecular components via the formation of membrane-bound and membrane-less compartments^1,2^. Membrane-less compartments, also known as biomolecular condensates^3,4^, are found both within the cytoplasm^5^ and nucleoplasm^6^. The underlying mechanism behind the emergence of biomolecular condensates is thought to be liquid-liquid phase separation (LLPS). LLPS broadly consists in the physicochemical demixing of biomolecules into different coexisting phases mediated by attractive interactions among multivalent biomolecules. In the cell environment, which is a highly multicomponent mixture of proteins, nucleic acids, and other molecules, the formation of a myriad of different condensates with diverse compositions is an intricate phenomenon. Phase separation is favoured when proteins, frequently containing intrinsically disordered regions (IDRs), and nucleic acids establish sufficiently favourable associative homotypic and/or heterotypic interactions with one another; that is, when such interactions result in a higher enthalpic gain, than those with the solvent, and that can compensate for the entropic loss due to demixing.

Biomolecular condensates display complex material properties spanning the range from low to high viscosity liquids, gels, and solids^7,8^. Furthermore, the material properties of condensates can change over time, for instance from liquid-like to solid-like, in a phenomenon described as ageing, rigidification, or maturation^9,10^. The liquid-to-solid transition in biomolecular condensates can be triggered by changes in the environmental conditions—such as in temperature^11^, ionic salt concentration^12^, pH or decreased adenosine triphosphate (ATP) levels^13^—that reduce protein solubility. Moreover, amino acid sequence mutations^14^, post-translational modifications^15^, or the application of mechanical forces are additional factors^7,16–^ that can contribute to the loss of the liquid-like character of a condensate over time (i.e., ageing^19,20^). These parameters are expected to, directly or indirectly, increase the proportion of long-lived protein contacts within the condensates, inducing complex mesoscale properties resulting from the rigidification of the molecular network within the condensate^9,21^.

The liquid-to-solid transition of condensates emerges from hindered dynamical behaviour of biomolecules inside condensates. Not surprisingly, such transition is implicated in the onset of several neurodegenerative and age-related diseases^10,22–25^. Certain mutations in the *fused in sarcoma* (FUS) protein^23,24^ and the TAR DNA-binding protein of 43 kDa (TDP-43)^14^, present in amyotrophic lateral sclerosis (ALS) patients, increase the speed and strength of condensate gelation^10^. Alike, mutations in *α*-synuclein—a protein associated with Parkinson’s disease^22^—and the Alzheimer-related *τ* -protein^26,27^ can induce LLPS and subsequent ageing into solid-like aggregates through the formation of *β*-amyloid fibrils.

The formation of inter-protein kinked *β*-sheets has been postulated as an intrinsic mechanism that can explain how condensates may undergo ageing without the need to invoke chemical changes in the system or in the environmental conditions^28^. Low-complexity domains (LCDs) with high aromatic content, which are important contributors to the multivalency of many naturally occurring proteins^4,29,30^, can exhibit local disorder-to-order structural transitions inside condensates, forming inter-protein *β*-sheets^14,28,31,32^ and, subsequently, amyloid fibrils^20,33,34^. The accumulation of inter-protein *β*-sheet structures formed by low-complexity aromatic-rich kinked segments (LARKS) produces concomitant enhancement of inter-protein interactions and can drive a gradual phase transition from functional liquid-like states into less dynamic reversible hydrogels or even irreversible solid-like states^7,10,35,36^. Indeed, the assembly of multiple LARKS into inter-locking fibril-like structures stabilized by *π*–*π* interactions and hydrogen bonding between backbone atoms can result into inter-peptide binding strengths of up to 50-75 *k*_B_*T* ^32,37,38^. Numerous naturally occurring phase-separating proteins such as FUS^28^, TDP-43^14,39^ or the heterogeneous nuclear ribonucleoprotein A1 (hnRNPA1)^7,40^, which are involved in the formation of the stress granules, are susceptible to LARKS– driven fibrillization within protein crowded environments (e.g., the interior of a biomolecular condensate). Hence, understanding the factors that promote protein fibrillization has become a key area of research, as it might contribute to devising strategies to prevent pathological transitions in condensates^41–45^.

In this work, we use multiscale simulations to investigate the impact of LARKS abundance and patterning in the progressive maturation of biomolecular condensates. Our multiscale approach bridges atomistic force fields^46,47^ and coarse-grained models of proteins and nucleic acids^48,49^. Based on structured *vs*. disordered peptide binding interactions predicted from free-energy atomistic calculations, we develop a coarse-grained model for LARKS-containing low-complexity domains that considers the accumulation of local structural transitions over time inside coarse-grained protein condensates. With this dynamical model, we unveil how the abundance and location of LARKS regions within the amino acid sequence impact the maturation rate (i.e., number of structural disorder-to-*β*-sheet transitions over time), protein network connectivity, density, and viscosity of condensates upon phase-separation. Importantly, we identify patternings that present moderate maturation rates, and enable condensates to maintain their functional liquid-like behaviour for longer timescales. Then, by exploiting the output from our coarse-grained condensate simulations, we investigate the mechanism of ageing of the protein FUS. FUS is an RNA-binding protein (RBP) that is involved in different biological tasks (such as transcription, DNA repair, or RNA biogenesis^50^) and is found in ALS-related aggregates^10,25,51^. By appropriately re-locating the LARKS domains within the FUS sequence, we observe how the ageing of the condensates is significantly decelerated. Furthermore, when RNA is added at moderate concentration to the FUS condensates with re-located LARKS, the formation of inter-protein *β*-sheets is suppressed. An RNA-blocking mechanism impedes LARKS structural transitions by engaging the bordering domains of the rearranged LARKS with neighbouring RNA strands that disrupt LARKS high-density fluctuations. Taken together, the work presented here sheds light on a key area of research aiming to design novel therapies to prevent age-related human diseases caused by condensate misregulation.

## II. RESULTS AND DISCUSSION

### A. LARKS abundance in low-complexity domains critically modulates condensate densification upon maturation

To elucidate the impact of the LARKS abundance and location in liquid-to-solid transitions of low-complexity domain sequences, we first develop a minimal coarse-grained model for LARKS-containing peptides. A key feature of this model is that we parameterize the binding strength among LARKS regions differently, depending on whether the LARKS are fully disordered or are forming inter-peptide *β*-sheets. We obtain the different inter-protein binding strengths from all-atom Potential of Mean-Force (PMF) simulations of interacting LARKS peptides. Specifically, to estimate the free energy of binding among proteins when they transition from disordered to forming inter-protein *β*-sheets, we perform all-atom Umbrella Sampling Molecular Dynamics (MD) simulations^52^ of systems containing six identical TDP-43 interacting LARKS peptides (_58_NFGAFS_63_^14^). Simulations are performed in explicit water and ions at room temperature and physiological salt concentration using the a99SB-*disp* force field^46^. By computing the PMF as a function of the center-of-mass (COM) distance between one reference peptide, which we dissociate from the other peptides (see further details of these calculations in the Supplementary Material), we determine the binding free energy difference between the structured (kinked *β*-sheet motif; PDB code 5WHN) and the fully disordered state of a TDP-43 LARKS aggregate (Fig. 1(a)).

**Figure 1:**
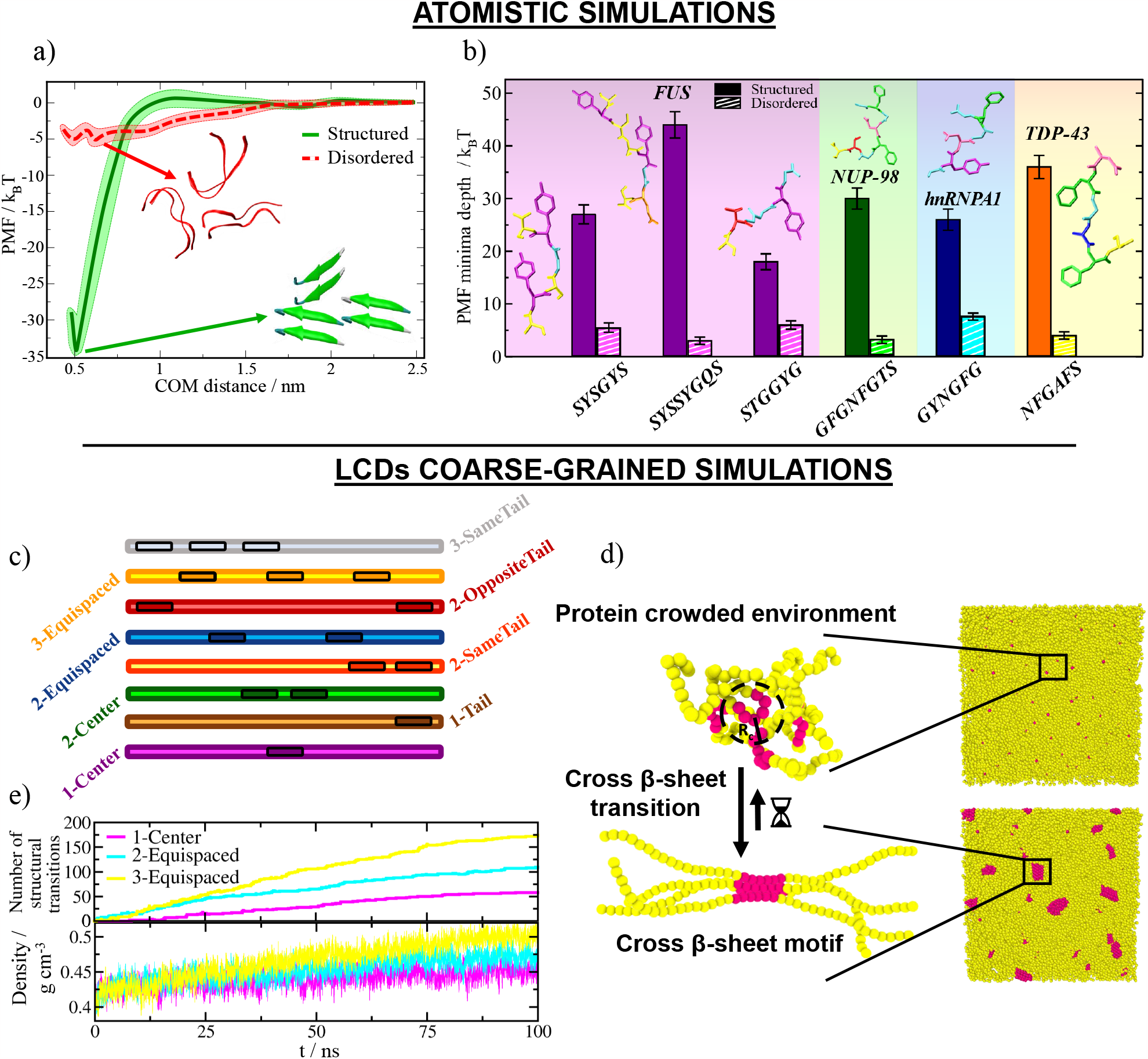
Disorder-to-*β*-sheet structural transitions augments protein binding, and their progressive accumulation within biomolecular condensates increases density. (a) Atomistic potential of mean force (PMF) dissociation curves of a 6-amino acid segment (PDB code: 5WHN) of the TDP-43 low-complexity domain sequence from a *β*-sheet structure formed by 6 peptides (of the same sequence) as a function of the center-of-mass distance (COM) using the a99SB-*disp* force field^46^. Simulations are carried out at room conditions and physiological NaCl concentration. While the solid green curve denotes the interaction strength among peptides with a kinked *β*-sheet structure, the dashed red curve accounts for the interaction energy among the same segments when they are fully disordered. Colour bands represent the statistical uncertainty of the PMF simulations. (b) Overview of the structured *vs*. disordered LARKS binding free energies found in the low-complexity domains of FUS^32^, NUP-98^53^, hnRNPA1^31^ and TDP-43 (this work). Snapshots of the atomistic LARKS sequences, included in the inset and specified below, are coloured by amino acid identity. (c) Schematic illustrations of the different low-complexity domain sequences proposed in this work to investigate LARKS abundance and patterning in condensate ageing. Susceptible regions to exhibit disorder-to-*β*-sheet transitions (i.e., 6-residue LARKS) are enclosed by black boxes. The total length of all low-complexity domain sequences is 125 residues. (d) Sketch of the ageing dynamic algorithm to mimic disorder-to-order transitions. Given a cut-off radius, if four LARKS segments encounter each other within such distance, an effective structural transition will be triggered and their binding strength is increased accordingly to the PMF free energy differences summarised in Panel (b). (e) Top: Number of structural transitions as a function of time for the three low-complexity domain sequences indicated in the legend. Bottom: Density time-evolution of the same sequences shown in the Top Panel.

Our atomistic simulations from Fig. 1(a) reveal that the inter-peptide binding energy is almost 7 times larger when the LARKS are forming *β*-sheets (solid green curve) than when they are fully disordered (dashed red curve). Such enhancement in binding free energy upon formation of *β*-sheets stems from the many hydrogen bonds that peptides at each step of the *β*-sheet ladder form with those in the next step of the ladder. Moreover, the additional stacking of aromatic side-chains that stabilize both the ladder and the individual *β*-sheets at each step confers global binding interaction strengths of ∼35 k_*B*_T per peptide (Fig. 1(a)). We note that the exact value of the difference in the free energy profile between the structured and disordered peptides is likely overestimated by the positional restraints, which enforce the stability of the *β*-sheet structures. Nevertheless, considering only the impact of constraints in our atomistic simulations is not sufficient to account for the observed enhancement of the inter-molecular protein interactions^32,53–55^. Such significant increase in the interaction energy upon the structural transition is consistent with the experimental observation of reversible hydrogels in several LARKS-containing RNA-binding proteins upon maturation (i.e., TDP-43 or FUS^14,56,57^), which can be dissolved with heat, and where a high percentage of *β*-sheet content has been found^11^. Furthermore, as reported in Fig. 1(b), the relatively high interaction strengths among structured LARKS that we obtain here for TDP-43 are also consistent with all-atom PMF calculations for NUP-98^53^, hnRNPA1^31^ and FUS^32^, previously reported by us. For all these proteins, a dramatic increase in binding energy from the disordered state (where interaction energies typically correspond to 3-8 k_*B*_T per peptide, suggesting that thermal fluctuations can easily break them) is observed upon the formation of the canonical *β*-sheet stacking (∼30-45 k_*B*_T of interaction per peptide). Alike, thermostable amyloid fibrils, such as those formed by the *Aβ*1–42, are expected to be stabilized by considerably high binding energies (i.e., of the order of 50-80 k_*B*_T)^37,38,58,59^. Overall, our atomistic simulations, presented in Figs. 1(a) and 1(b), provide quantitative information of the strengthening of inter-protein binding upon LARKS structural transitions for a set of RNAbinding proteins that exhibit progressive aberrant solidification^25,60^.

Based on our all-atom simulation results presented in Figs. 1(a-b), we now develop a coarse-grained model for low-complexity domain peptides, where we enforce *β*-sheet transitions through an algorithm which recapitulates the non-equilibrium process of condensate ageing due to strengthening of LARKS–LARKS interactions. Our model approximates protein low-complexity domains as fully flexible (pseudo) Lennard-Jones heteropolymers with beads connected by harmonic springs, in which each bead represents a single residue. The Wang-Frenkel potential^61^ is employed to describe the excluded volume and inter-molecular interactions between non-bonded residues of the low-complexity domains. Furthermore, to speed up simulations, we employ an implicit-solvent model in which the diluted phase (i.e., the protein-poor liquid phase) and the condensed phase (i.e., the protein-rich liquid phase) are effectively a vapor and a liquid phase, respectively. Within a single low-complexity domain, we combine regions prone to exhibiting disorder-to-*β*-sheet transitions (i.e., LARKS) with regions that can only establish weak and transient interactions. Importantly, we distinguish two main types of interactions: (1) weak interactions for disorder-like sticky amino acids (which are of the order of 0.5 k_*B*_T and can be easily overcome by thermal fluctuations as characteristic of liquid-like behaviour); and (2) strengthened *β*-sheet-like interactions (with a binding energy of approximately 3 k_*B*_T per amino acid and characteristic strength of gel-like behaviour). This energy-scale has been derived from the PMF results shown in Fig. 1(b) for FUS, NUP-98, hnRNPA1 and TDP-43 proteins (for further details on the model parametrization see the Supplementary Material). Therefore, along the low-complexity domains, we define regions that are susceptible to undergo a disorder-to-*β*-sheet transition (henceforth called ‘LARKS unstructured region’; dark highlighted domains in Fig. 1(c)), and domains which cannot exhibit a structural transition (henceforth called ‘non-LARKS unstructured region’; brighter domains in Fig. 1(c)). Then, through our dynamic algorithm^31,32^, we approximate the non-equilibrium process of condensate ageing by considering the atomistic implications (i.e., non-conservative strengthening of inter-protein binding, local protein rigidification, and changes in the inter-molecular organization; Fig. 1(a)) of the gradual and irreversible accumulation of inter-protein *β*-sheet structures in a time-dependent manner, and as a function of the local protein density. Whereas all low-complexity domains initially interact through weak and transient interactions (both LARKS and non-LARKS unstructured regions), the dynamic algorithm triggers transitions from unstructured LARKS to kinked *β*-sheets when one of the central beads of a given LARKS is in close contact (within a cut-off distance of ∼8 Å) with three other LARKS of neighbouring low-complexity domains^14,28,56^.

Therefore, dynamically, our algorithm evaluates whether the conditions around each unstructured LARKS are favorable for undergoing an ‘effective’ structural transition. If the conditions are met, the transition is recapitulated by enhancing the interaction strength and rigidity of the four LARKS involved in the *β*-sheet motif by a factor of 6—as observed through our atomistic PMF simulations for the studied set of RNA-binding proteins (Fig. 1(b)). This value corresponds to the ratio between structured and disordered binding energies averaged over all the studied sequences presented in Fig. 1(b). The comprehensive description of the coarse-grained potentials, a full list of the model parameters, as well as the studied low-complexity domain sequences (shown in Fig. 1(d)) and details on the dynamic algorithm are provided in the Supplementary Material.

With this coarse-grained model for low-complexity domain peptides and our dynamical algorithm, we investigate the impact of LARKS abundance in condensate maturation. Since most of the known low-complexity domain sequences which display progressive condensate rigidifcation contain at most three LARKS^14,28,56^, we compare the maturation rate (i.e., number of structural transitions over time) of condensates made of peptides containing 1, 2 or 3 LARKS. The chosen LARKS distribution across the low-complexity domains is in all cases symmetric and equispaced (as shown in Fig. 1(c) for the ‘1-Center’, ‘2-Equispaced’ and ‘3-Equispaced’ sequences) and the total length of the low-complexity domains is set to 125 amino acids—a typical length of a low-complexity domain in a phase-separating RBP^33,62^. Moreover, the peptide length of each LARKS corresponds to six amino acids (Fig. 1(b)), and their interactions are initially disorder-like (i.e., ∼0.5 k_*B*_T per residue; as those of the rest of the low-complexity domains) but susceptible to undergo a structural transition over time (Fig. 1(d)). Importantly, in low-complexity domains with more than one LARKS, we limit inter-peptide *β*-sheet transitions to occur only within clusters of peptides of the same identity—i.e., 1-1, 2-2 or 3-3—to avoid autoinhibited low-complexity domain states^63^, and to mimic LARKS sequence specificity^28,64^. Using Direct Coexistence simulations^65,66^, we estimate the phase diagrams prior to ageing for solutions of the three different low-complexity domain peptides in the temperature-density (*T* − *ρ*) plane (Fig. 2(a) black curve; further details on these calculations are provided in the Supplementary Material). By construction, the three phase diagrams are equivalent prior to ageing, but they become significantly different post-ageing. The ageing behaviour is investigated below the critical temperature (*T/T*_*c*_=0.91) by means of bulk *NpT* simulations (starting from the coexisting condensed density pre-ageing) with our ageing dynamic algorithm that implements structural transitions in a local and time-dependent manner (Fig. 1(d)). As expected, the accumulation of *β*-sheet content over time is significantly higher for proteins with more LARKS domains (Fig. 1(e) Top panel). Moreover, ageing of protein condensates with more numerous LARKS also results in significant enhancement of their density over time (Fig. 1(e) Bottom panel). That is, within the same timescale, aged condensates formed by proteins containing a single LARKS increase their density by a 10%, and those made of proteins with three LARKS present densities 25% higher. These results can be explained by the larger number of high-density packed domains (e.g., cross-*β*-sheets) found in condensates formed by proteins with a higher number of LARKS (Fig. 2(f)). Furthermore, these results are consistent with recent experimental observations of FUS-LCD droplets aged via thermal-annealing, where a rise in droplet density correlated with an increase in *β*sheet content^11^.

**Figure 2:**
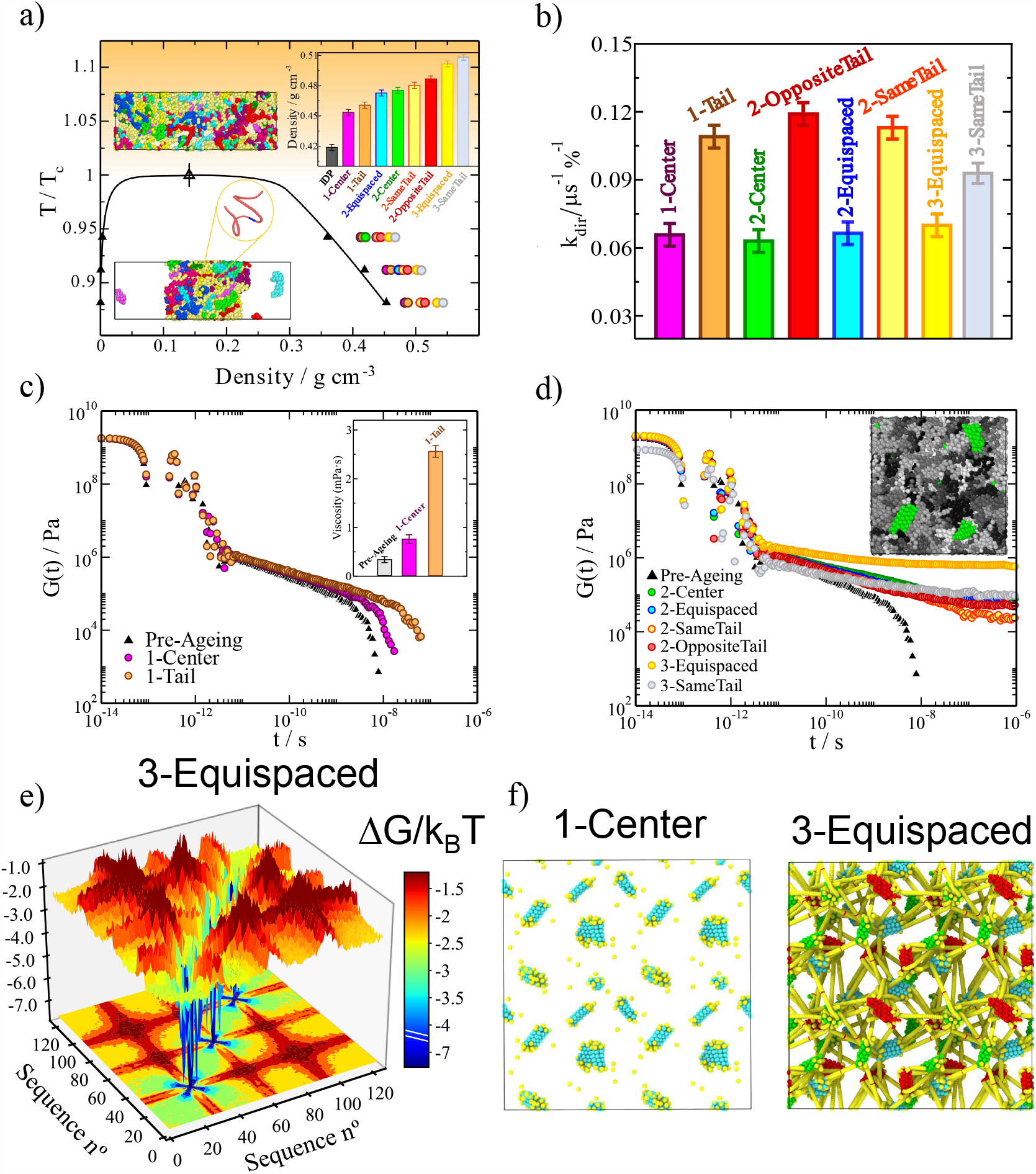
LARKS abundance and patterning in low-complexity domains strongly determine the liquid network connectivity and viscoelastic properties of aged condensates. a) Phase diagram in the *T* − *ρ* plane for all different low-complexity domain sequences prior ageing (black curve). Coloured symbols indicate the condensate density of the distinct low-complexity domains upon maturation as emphasized in the inset for *T/T*_*c*_=0.91. The orange background indicates temperatures at which LLPS is not observable (i.e., *T/T*_*c*_ *>* 1). Snapshots of a Direct Coexistence simulation of the ‘1-Center’ sequence (zoom-in) exhibiting LLPS (Bottom) and non-demixing behaviour (Top) are also included. Each low-complexity domain replica is depicted in a different colour. b) Kinetic constants (in *μs*^−1^·%^−1^) of inter-protein *β*-sheet formation for the different low-complexity domain sequences shown in Fig. 1(c) and Panel (a). c-d) Shear stress relaxation modulus *G*(*t*) for low-complexity domain phase-separated condensates pre-ageing (black triangles) and post-ageing (coloured circles) as indicated in the legend of each panel. Please note that by construction, all low-complexity domain sequences possess the same viscoelastic behaviour and phase diagram before maturation. The inset in Panel (c) shows the condensate viscosity pre-ageing and post-ageing for the ‘1-Center’ and ‘1-Tail’ sequences. The maturation time after which all these calculations were performed was 0.1 *μs*. In panel (d) a snapshot of an aged bulk condensed phase is shown: *β*-sheets clusters are coloured in green while low-complexity domain unstructured regions in grey tones (where different tones are used for distinct low-complexity domain replicas). e) Landscape of the protein contact free energy variation upon condensate maturation for the ‘3-Equispaced’ low-complexity domain sequence. ΔG/k_*B*_T is obtained from the residue contact probability ratio between aged condensates and liquid-like condensates before ageing. A colour map projection of the free energy landscape in 2-dimensions is also included. f) Network connectivity of aged ‘1-Center’ and ‘3-Equispaced’ low-complexity domain condensates evaluated using the PPA method.

### B. LARKS located at the ends of the low-complexity domains promote faster ageing than those located at the center

We now investigate the role of the location of the LARKS sequences within a protein on the ageing of its condensates. To that aim, we design eight distinct low-complexity domains containing either one or two LARKS domains but located on different regions of the protein, as shown in Fig. 1(c) and detailed in the Supplementary Material. Specifically, we consider the following proteins: with one LARKS in the center (‘1-Center’), one LARKS at the end of the protein or tail (‘1-Tail’), two LARKS in the center (‘2-Center’), two LARKS near the same tail (‘2-SameTail’), one LARKS each on each tail (‘2-OppositeTail’), and two LARKS equispaced across the whole protein (‘2-Equispaced’). Moreover, to distinguish between the effects of LARKS location and abundance, we also investigate the behaviour of a three-LARKS equally-distributed low-complexity domain sequence (‘3-Equispaced’) and three LARKS near the same tail (‘3-SameTail’). In Figure 2(a), we show the phase diagrams (in the *T* − *ρ* plane) for these different proteins pre-ageing (black curve; please note that all low-complexity domains before maturation possess the same phase diagram) as well as the post-ageing condensate densities at different temperatures. Importantly, independently of the number of LARKS or their patterning, in all cases, the continuous accumulation of *β*-sheets augments the density over time—due to the higher packing fraction of these structured motifs with respect to clusters of disordered domains—until condensates become kinetically arrested. Nevertheless, neither the patterning nor the abundance of LARKS, alter the critical temperature (*T*_*c*_) below which phase separation is observed. That is because high-density fluctuations driving *β*-sheet transitions can only take place in phase-separated protein liquid phases^67^, and therefore, the critical conditions (i.e., temperature or saturation concentration) for observing LLPS remain unaffected. Strikingly, we find that proteins with tail-located LARKS exhibit higher increments in condensate density than proteins with center-located and equispaced LARKS. Such behaviour occurs for proteins with 1-LARKS, 2-LARKS and 3-LARKS (Figure 2(a): inset); reaching the ‘2-OppositeTail’ patterning slightly higher droplet density than the ‘2-SameTail’ sequence. Furthermore, LARKS abundance is also critical for controlling droplet densification upon maturation, where the relative increase in density is 2.5 times higher from 1-LARKS to 3-LARKS low-complexity domains. These results are consistent with previous *in silico* observations for FUS-LCD and A1-LCD-hnRNPA1 aged condensates, where the higher concentration of LARKS within the FUS-LCD sequence led to higher densification upon maturation compared to A1-LCD-hnRNPA1 droplets^31^. Recently, FUS-LCD ageing-driven densification through *β*-sheet aggregation has been confirmed^11,67^.

Next, we quantify the maturation rate of the different condensates (Fig. 1(c)) by using a simple kinetic chemical model to fit the time-evolution of the number of structural transitions found in our simulations. The model used includes two forward reactions: 2[*D*] → 2[*S*] and [*D*] + [*S*] → 2[*S*]; and one backward reaction [*S*] → [*D*], where [*D*] represents the percentage of fully disordered LARKS, and [*S*] the percentage of LARKS forming inter-protein *β*-sheets. Such analysis allows us to estimate the kinetic constant *k*_*dir*_ of interprotein *β*-sheet formation (further details are provided in the Supplementary Material). Remarkably, the kinetic constants obtained for all proteins investigated with LARKS positioned on the tails (i.e., ‘1-Tail’, ‘2-OppositeTail’ and ‘2-SameTail’) roughly double the *k*_*dir*_ value of those in which LARKS are center-located or equidistributed along the sequence (Figure 2(b)). Such behaviour emerges because, at short times, the ends of the proteins explore a larger volume of the condensate than the segments located in the central part^68^; thus, enhancing the probability of the tails to sample high-density protein fluctuations within the condensates. Furthermore, polymer termini typically exhibit a higher ratio of inter/intra molecular contacts than central regions within liquid phases^69^. The only apparent exception is the ‘3-SameTail’ LCD, which exhibits intermediate ageing kinetics between tail and center-located sequences. However, this is a special case because to keep constant the minimum LARKS interspacing imposed in the 2-SameTail LCD (e.g., 14 residues) one of the three LARKS in the 3-SameTail LCD needs to be positioned near the center of the sequence (Fig. 1(c)). Accordingly, among all the studied sequences, the ‘2-OppositeTail’ presents the highest structural transition *k*_*dir*_, whereas the ‘2-Center’ the lowest value, being twice slower its maturation rate compared to the former. Overall, our results from Fig. 2(c) suggest that, independently of the number of LARKS, LCDs that contain LARKS positioned towards their end regions should age faster.

The different rates of structural transitions (Figure 2(b)) led us to investigate the relationship between *β*-sheet content and the viscosity of the various aged condensates. The viscoelastic properties of a system can be measured by the time-dependent mechanical response of the material in bulk conditions when it is subjected to a small shear deformation. That is described by the shear stress relaxation modulus *G*(*t*). In principle, *G*(*t*) can be computed from the auto-correlation of any of the off-diagonal components of the pressure tensor^70^. However, in practise, the statistical uncertainty is significantly reduced by considering all the elements of the pressure tensor^71,72^. Then, the zero-shear-rate viscosity (*η*) of the material is obtained by integrating with respect to time the stress relaxation modulus^71^ (see Supplementary Material for further details on these calculations). We evaluate *η* by means of bulk NVT simulations in which the density of the system corresponds to that of the condensate in bulk conditions—hence approximating that both within the condensate’s interface and its core, the rate of cross-*β*-sheet transitions is of the same order^17,32,64^. In Fig. 2(c), we compare *G*(*t*) for the pre-ageing (black triangles) and post-ageing ‘1-Center’ (purple circles) and ‘1-Tail’ (orange circles) proteins. We observe that the decay of the shear stress relaxation modulus occurs at much longer timescales for both aged condensates, especially for that composed of ‘1-Tail’ low-complexity domain sequences. Such slower decay in *G*(*t*) translates into condensate viscosities upon maturation that are almost 10 times higher than those prior to ageing, as it occurs for the ‘1-Tail’ sequence (Fig. 2(c) inset). Indeed, the LARKS location within the protein plays a critical role in droplet maturation, being the viscosity for ‘1-Tail’ condensates more than 3 times higher than that for ‘1-Center’ droplets. These results, on top of those shown in Figs. 2(a) and 2(b) for progressive droplet densification and the structural transition rate respectively, emphasise the relevance of LARKS location in determining the characteristics of condensates during and post-ageing.

If we now examine the viscosity increase for the different 2-LARKS and 3-LARKS proteins upon the same ageing timescale that we explored for the 1-LARKS sequences in Fig. 2(c), we observe that none of the aged condensates presents a decay in *G*(*t*) (Fig. 2(d)). Instead, the shear stress moduli fall into persistent plateaus with no hints of decaying at comparable timescales, and yielding to infinite viscosity values (and non-diffusive behaviour) characteristic of gel-like states. Such type of behaviour was recently observed experimentally for aged FUS condensates^9^. These results strongly suggest that low-complexity domains with at least two LARKS, independently of their patterning, can progressively form gel-like or solid-like aggregates through *β*-sheet fibrilization. Nevertheless, proteins with a single LARKS, despite gradually increasing their viscosity over time, may not attain complete kinetic arrest due to the impossibility of forming a 3-dimensional (*β*-sheet) percolated network characteristic of a gel-like state^31^. Instead, they may exhibit persistent high viscuous liquid-like behaviour with extremely long relaxation timescales due to a progressive slowdown in protein mobility, as recently reported for A1-LCD-hnRNPA1 condensates^7,40^. Interestingly, since the evaluation of *G*(*t*) not only provides fundamental information on how the material properties of condensates change during ageing, but also on how such changes are dictated by different relaxation mechanisms, we find that the shear moduli for all proteins studied are almost identical at short timescales (Figs. 2(c) and 2(d)). Consequently, the timescales of fast events, such as formation and breakage of short-range interactions and of intramolecular reorganisation (i.e., internal protein conformational fluctuations, such as bond or angle relaxation modes) of non-aged and aged condensates are expected to be remarkably similar in the studied condensates. However, at long timescales, where the stress relaxation modulus is mainly dominated by intermolecular forces, long-range conformational changes (i.e. protein folding/unfolding events), and protein diffusion within the crowded condensate environment, the effect of LARKS patterning and abundance is decisive in determining the viscoelastic properties of the droplets. Consistent with our results from Figs. 2(c) and 2(d), a deceleration in protein mobility during ageing has been experimentally measured through higher condensate viscosities^22,73^, and lower or incomplete Fluorescence Recovery after Photobleaching (FRAP) measurements^22,74–79^. Furthermore, techniques such as GFP fluorescence recovery, coalescence, and active and passive microrheology experiments have revealed how over time liquid-like condensates formed by proteins with low-complexity domains, especially those with high aromatic content, can transition to gels or soft glasses^9,10,76,80^. Several factors have been proposed as key drivers for the liquidto-solid transition, including altered salt-concentration or temperature^9,81^, post-translational modifications^26,82^ and protein mutations^16,83,84^. In particular, the accumulation of *β*-sheet transitions has been put forward as an important mechanism that triggers condensate ageing by increasing the binding affinity among species and slowing down the timescales of inter-protein unbinding events^7,28,36,67^. Our results presented here provide strong evidence of the critical importance of an additional parameter: LARKS concentration and location.

### C. Cross-*β*-sheet percolated networks are hallmarks of aberrant condensation

We next characterise, at molecular resolution, the changes in structure, topology and free energy of the intermolecular network of interactions in the condensates as a result of ageing. For this, we perform an energy-scaled molecular interaction analysis, which has been recently proposed in Refs.^85,86^. In such analysis, the probability that any two residues of the different proteins inside the condensate can contact one another is estimated through a cut-off distance analysis that considers the residue mean excluded volume and the minimum potential energy of each interacting pair. From the difference in the energy-scaled contact probability between pre-aged (*P*_*pre*_) and aged condensates (*P*_*post*_), we can estimate the free energy difference of interactions 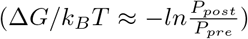, which relates to the progressive transformation of the liquid-network connectivity as a result of the bond strengthening triggered by ageing (details on these calculations can be found in the Supplementary Material). Hence, through this analysis, we can unveil the microscopic pathway by which proteins containing more than one LARKS transition from liquid-like behaviour to gel-like kinetically trapped states, as shown through the stress relaxation modulus in Fig. 2(d). Furthermore, these results are supported by additional calculations of the shear storage (G’) and shear loss (G”) moduli (Fig. S8), showing that the presence of two or more LARKS along the sequence lead to higher values of G’ with respect to G”, and thus, suggesting gel-like behaviour.

The condensate connectivity free energy difference upon maturation shown in Fig. 2(e) for the ‘3-Equispaced’ sequence reveals large increments in binding energy (of up 7 k_*B*_T per amino acid) driven by LARKS *β*-sheet transitions. These values are fully consistent with those obtained by atomistic simulations for TDP-43, FUS, hnRNPA1, and NUP-98 (Fig. 1(b)), where the binding free energy differences between ordered and disordered states ranged from 3-8 k_*B*_T per amino acid depending on the considered LARKS sequence. Interestingly, we also observe that LARKS adjacent domains significantly enhance their binding interaction by 1 to 3 k_*B*_T due to neighbouring crossed-*β*-sheet accumulation and the subsequent droplet densification. Nevertheless, we note that due to the non-sequence specificity of our protein model (e.g., not considering long-range repulsion between residues of the same charge), the values for LARKS adjacent binding strengthening might be slightly overestimated^31^. Overall, Fig. 2(e) energetically explains the observed plateau in *G*(*t*) after condensate maturation (Fig. 2(d)) for the 3-LARKS sequence. The connectivity free energy difference upon condensate ageing for the rest of protein sequences is shown in Fig. S5. Notably, while the local strengthening of bonds is of the same order (i.e., 6-7 k_*B*_T) for all protein sequences, the abundance and location of LARKS has a crucial importance in the maturation rate and viscoelastic properties of the aged droplets.

Another key parameter that describes the microscopic behaviour of aged condensates is the structural features of the intermolecular network of protein molecular connections^53^. To characterise the topology of such a network, we employ a modification^31^ to the primitive path analysis (PPA) method originally developed to study the structure of polymer melts^87,88^. Within this approach (further described in the Supplementary Material), we consider that inter-peptide *β*-sheet bonds are fixed in space, that the intra-molecular excluded volume is zero, and that the bond length has an equilibrium value of 0 nm. These considerations allow us to neglect the contour length of protein strands that connect different LARKS, while preserving the topology of the underlying network. Then, at the end of the minimization, we can visualize the network of elastically inter-protein *β*-sheet clusters that determines the aged condensate viscoelastic behaviour. In Fig. 2(f), we show the *β*-sheet percolation network for the ‘1-Center’ (Left) and ‘3-Equispaced’ (Right) low-complexity domain aged condensates upon the same maturation timescale (0.1 *μs*; please note that the absence of explicit solvent significantly accelerate the dynamics of our model). Our results reveal a remarkable difference in the molecular connectivity between ‘1-Center’ and ‘3-Equispaced’ aged condensates. Consistent with the vis-coelastic properties of the 3-LARKS sequence, in which aged condensates display gel-like behaviour (Fig. 2(d)), we observe a network of strong inter-protein *β*-sheet contacts which fully percolates throughout the condensate. Such tightly connected network is precisely what inhibits the relaxation of proteins within the droplets at long timescales and explains the observed rubbery plateau in the stress relaxation modulus suggesting gel-like behaviour. In contrast, the inter-protein *β*-sheet network in aged ‘1-Center’ low-complexity domain condensates presents only disconnected gel-like domains. In this case, the lack of percolation of the *β*-sheet network allows the condensate to relax as a whole at long timescales, as dictated by the decay to zero of *G*(*t*) (Fig. 2(c)). Nevertheless, still 1-LARKS aged condensates exhibit much higher viscosities than non-aged droplets which are weakly sustained by short-lived protein binding^63,89^. Hence, according to our results, shown in Figs. 2(d) and 2(f), only proteins with at least 2 strong-binding anchoring points (which can bind to multiple LARKS at the same time) can establish the percolated network characteristic of a non-diffusive gel^70^. Furthermore, the progressive dynamical arrest through the emergence of imbalanced inter-molecular interactions (Fig. 2(e)) giving rise to *β*-sheet percolated networks widely explains the recognised asphericity of aged condensates^9,22^ and the emergence of irregular morphologies through non-ergodic droplet coalescence^11,53^, reported for proteins containing low complexity domains, such as hnRNPA1^20^, FUS^9^, TDP-43^90^, or NUP-98^91^.

### D. Sequence domain reordering of FUS decelerates *β*-sheet accumulation in aged condensates

We now aim to exploit the insight obtained in the previous sections to elucidate possible routes to inhibit, or at least decelerate, the progressive rigidifcation of FUS condensates. We have chosen FUS because it is an RNA-binding protein, whose phase behaviour^33,62,92^ and ageing^10,23–25,51^ have been widely investigated. FUS-containing condensates are involved in the formation of stress granules^93^, transcription^94^, DNA repair and RNA biogenesis^95^, and their ageing have been implicated in several neurodegenerative disorders such as ALS and frontotemporal dementia (FTD)^25,60^. Importantly, the main driving force causing gradual solidification of FUS in absence of external stimuli, or changes in the thermodynamic conditions, has been suggested to be the accumulation of inter-protein *β*-sheets^28,67,96^. FUS is a multidomain protein (LCD, RGG1, RRM, RGG2, ZF, and RGG3; see Fig. 3(a) Top). Thus, to try to perturb the formation of inter-protein *β*-sheets at the LARKS of FUS inside the condensates, we reposition the different domains of FUS (Fig. 3(a) Bottom). Specifically, we place the LARKS-containing LCD in the middle of the FUS sequence. According to our results shown in Figs. 2(b) and 2(c), this approach should decrease the probability of the LARKS to form part of a high-density protein cluster, hence, decelerating the *β*-sheet transition rate within condensates and its associated increase of viscosity over time. Moreover, the entropy of the condensed phase should be lower when long-lived *β*-sheet pairings are closer to the center of the chain than to its tails^97^. We reasoned that in case that FUS phase-separation would occur in presence of RNA^18,98,99^, such reordering could also potentially frustrate LARKS high-density fluctuations by virtue of the FUS arginine-glycine rich-regions (RGGs) and RNA-recognition motifs (RRMs) flanking the low-complexity domain (Fig. 3(a)) and recruiting high concentrations of RNA^62,85^. In addition to the reordering of domains, we also test the impact of mutating adjacent glycines (G) and serines (S) to the three FUS LARKS by arginines (R) across the original FUS sequence (Fig. 3(a) Middle). We hypothesize that such mutations may disrupt *β*-sheet binding through repulsive electrostatic interactions (R-R) or even favourable unstructured LARKS-nonLARKS cation-*π* interactions; and it could facilitate RNA-LCD binding (due to the higher content of arginines within the LCD, instead of LLPS spacers such as glycines or serines^100^; Fig. 3(a) Middle) precluding the LCD high-density fluctuations triggering structural transitions^31^.

**Figure 3:**
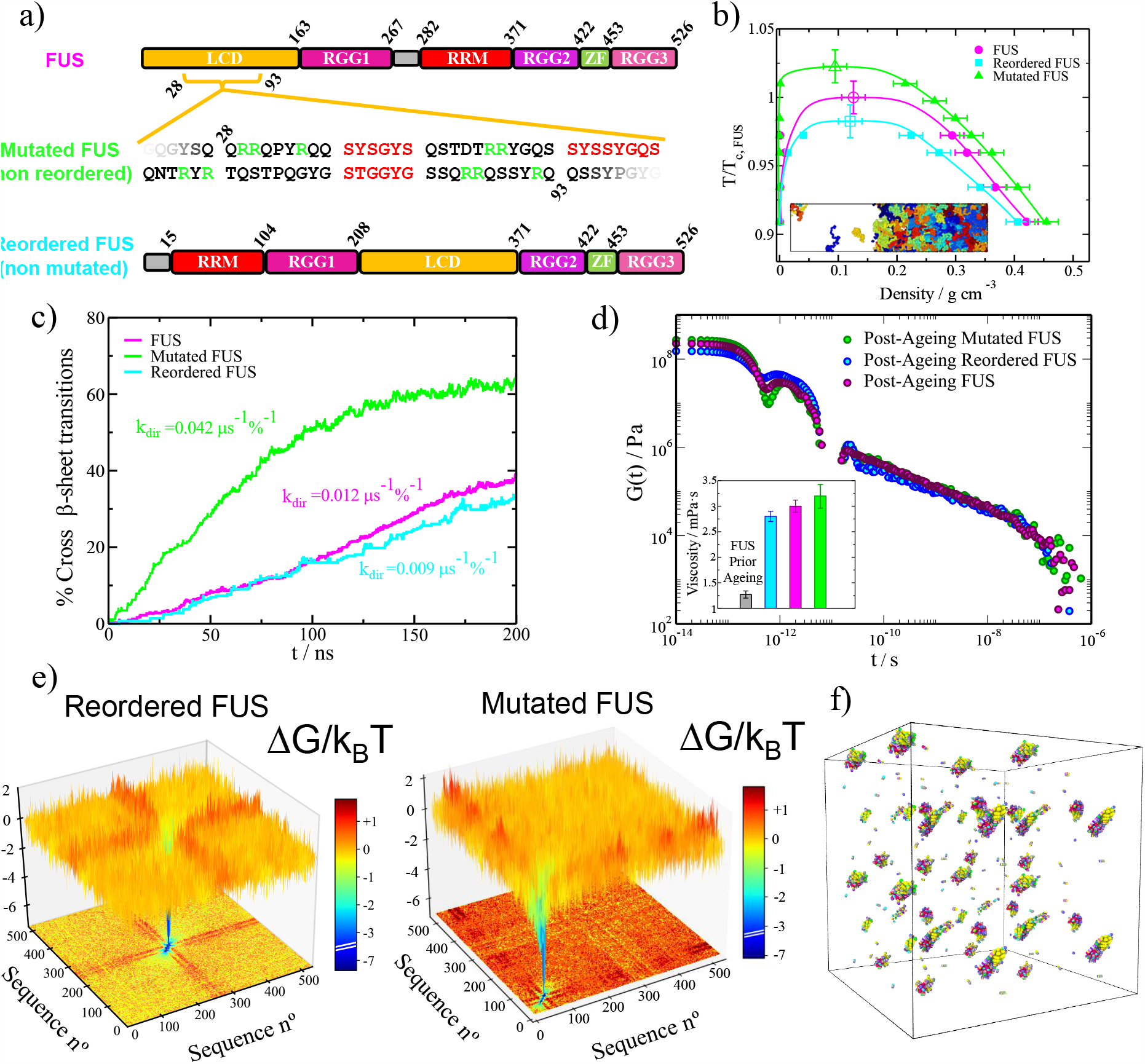
Sequence domain reordering in FUS decelerates the accumulation of crossed-*β*-sheet motifs in phase-separated condensates. a) Different domains of the human FUS sequence (Top) and the reordered FUS variant (Bottom) where LCD accounts for low-complexity domain, RGG1, RGG2 and RGG3 for the different Arginine-Glycine-rich regions (RGG), RRM for the RNA-recognition motif, and ZF for the zinc finger. Please note that the only difference between these two sequences is the location of their distinct domains. In the Mid Panel, we include the performed mutations (serines and glycines by arginines; green) on the adjacent regions of the three LARKS (depicted in red) of the low-complexity domain in FUS. Please note that the mutated FUS sequence contains the same domain ordering as the original sequence. b) Phase diagram in the *T* − *ρ* plane for the three studied FUS sequences as indicated in the legend. Temperature is renormalized by the critical temperature of pure FUS (*T*_*c,F US*_). A snapshot of a Direct Coexistence simulation of FUS is also included with each protein replica coloured with a different tone. c) Time-evolution of the percentage of LARKS forming crossed-*β*-sheet motifs in phase-separated condensates of the three FUS variants at *T/T*_*c,pure*_ ∼0.97 (referring *T*_*c,pure*_ to the critical *T* of each FUS sequence). The kinetic constant (*k*_*dir*_) evaluated through a second-order reaction analysis is provided for each curve. d) Shear stress relaxation modulus *G*(*t*) of the different FUS condensates (at *T/T*_*c,pure*_ ∼0.97) after 0.3 *μs* of maturation. The viscosity for each FUS condensate after maturation is included (coloured bars) along the pre-ageing viscosity for the original FUS sequence (grey bar) in the inset. e) Landscape of the protein contact free energy variation upon condensate maturation for the FUS reordered (Left) and FUS mutated variants (Right). Colour map projections of the free energy landscape in 2-dimensions are also included. f) *β*-sheet network connectivity of an aged reordered FUS condensate evaluated through the PPA method.

To perform these simulations, we use a sequence-dependent residue-resolution force field (HPS-Cation-*π*^101,102^) which qualitatively reproduces the propensity of numerous RNA-binding proteins (including FUS) to phase separate at physiological conditions^85^, as well as their RNA-concentration-dependent reentrant phase behaviour^92,103–105^. Furthermore, to enable *β*-sheet binding strengthening (as in our previous low-complexity domain simulations), we employ a dynamic algorithm that mimics transitions from disordered LARKS to interprotein *β*-sheets when the central C_*α*_ residue of a LARKS is in close contact (within a given cut-off distance: ∼8 Å) with three other LARKS of neighbouring proteins^14,28,56^. Such structural transitions are enforced by enhancing the interaction strength and rigidity of the involved LARKS (as in our low-complexity domain simulations shown in Figs. 1 and 2). The corresponding binding strengthening among each FUS-LARKS sequence has been derived from the atomistic PMF simulations shown in Fig. 1(b). Further details on the HPS-Cation-*π*^101,102^ force field, dynamical algorithm, and the local order parameter driving structural transitions are described in the Supplementary Material. In Fig. 3(a), we summarize the three FUS sequences for which we investigate condensate maturation. In the upper part of the panel, we show the original full-FUS sequence broken down by domains. In the mid panel, we describe the performed mutations in the low-complexity domain of the original FUS protein (indicated in green), where also the three LARKS domains are highlighted in red. Finally, in the lower panel, we show the applied low-complexity domain reordering (with no mutations) to the original FUS sequence, in which the low-complexity domain is relocated in between the RGG1 and RGG2 regions. The phase diagram prior to ageing (in the *T* − *ρ* plane) for the standard FUS, the reordered FUS, and the mutated FUS systems are reported in Fig. 3(b). Remarkably, in the pre-ageing regime, the estimated critical temperature of both FUS variants differ in less than 2.5% with the original sequence, hence, demonstrating their similar ability to phase separate.

In vitro experiments can now routinely investigate the phase behaviour of proteins with specific amino acid mutations. For instance, the phase diagrams of 28 A1-LCD variants have been quantified experimentally^100^ and the impact of amino acid mutations on the phase behaviour of the FUS protein have been investigated^106^. It is worth noting that the FUS variants proposed in this work (Fig. 3(a)), where the LARKS domains are reordered, may be challenging to produce experimentally and are not present in cells. Nonetheless, protein *ad hoc* sequence modifications can be achieved experimentally. For instance, the RGG domain of LAF-1 has been reshuffled experimentally, inducing a significant enhancement in the protein’s phase separation propensity compared to the unmodified variant^107^. IDPs can be also be synthesized with *ad hoc* aromatic content and amino acid sequences^108^.

Next, we investigate the progressive accumulation of inter-protein *β*-sheets inside the phase-separated condensates of these three different proteins. To that goal, we set condensed phase conditions at the corresponding coexisting density of each FUS protein at *T/T*_*c,pure*_ ∼0.97 (referring *T*_*c,pure*_ to the critical temperature of each FUS sequence), and we measure the number of LARKS structural transitions (in percentage) as a function of time (Fig. 3(c)). Then, through the same second-order reaction analysis used in Fig. 2(b), we evaluate the kinetic constant of *β*-sheet formation upon maturation. Notably, we find that *k*_*dir*_ is significantly lower (25%) for the reordered-LCD FUS variant (cyan curve) when compared to the original FUS protein (magenta curve). These results further support our previous findings with different minimal peptides, in which lower kinetic constants were found consistently for proteins with LARKS located towards the middle of the sequence (Fig. 2(b)). On the other hand, the FUS mutated arginine-rich variant (green curve) presents a substantially higher *k*_*dir*_ (of roughly 300%) than the original sequence (Fig. 3(c)). Interestingly, the overall balance between favourable interactions (among arginine and aromatic residues) *vs*. unfavourable electrostatic repulsion (between positively charged arginine-rich patches) leads in this case to an increase of high-density fluctuations of LARKS rich-regions. That directly translates into a severely higher accumulation of crossed-*β*-sheet fibrils over time (Fig. 3(c)). Interestingly, in many proteins—including RBPs such as FUS, TDP-43 or hnRNPA1—the LCD is preferentially located at the extremities of the protein^109^. This might be an evolutionary aspect to maximize the degree of inter-protein connectivity^109^. Thus, repositioning the LCD (where LARKS are usually found) towards the center of the sequence should reduce inter-protein molecular binding and concomitant *β*-sheet accumulation. We have chosen hnRNPA1 as a work-case example of a single LARKS-containing protein^28^ to obtain more general conclusions about this domain-reordering approach. We have designed a reordered hnRNPA1 variant (see Fig. S9(a)) in which we place the LCD in the center of the protein. The pre-ageing phase diagrams of both hnRNPA1 and the LCD-reordered variant differ in less than 2% on their critical temperatures (Fig. S9(b); similar to what we have found for FUS wild-type and their two variants in Fig. 3(b)). Now, if we determine the inter-protein *β*-sheet transition rate within phase-separated condensates of both variants, we find that *k*_*dir*_ for the reordered hn-RNPA1 variant is approximately half the kinetic constant of the wild-type hnRNPA1 sequence (Fig. S9(c)). Thus, taken together, our findings for hnRNPA1 and FUS reveal a general behaviour: the reallocation of the protein LARKS towards the center of the sequence decelerates the accumulation of inter-protein *β*-sheets.

To characterise the post-ageing droplet viscoelastic behaviour of the different FUS proteins, we measure the shear stress relaxation modulus after 0.3 *μs* of maturation time (Fig. 3(d); we note that the coarse-grained nature of our model including implicit-solvent^101,102,110^ overestimates the kinetics of protein diffusion in our simulations^85^). By integrating in time *G*(*t*), we reveal that the reordered-LCD FUS variant possesses the lowest viscosity when compared to the original and mutated proteins after ageing. In contrast, the mutated arginine-rich variant presents the highest viscosity, which is consistent with its higher emergence rate of crossed-*β*-sheet clusters (Fig. 3(c)). Notwithstanding, the viscosities of the three sequences after ageing are almost 3 times higher than those of the original FUS protein prior to ageing (Fig. 3(d) inset).

To understand from a molecular perspective the gradual transformation of the different variants during ageing, we now evaluate the change to the free energy of the intermolecular interaction network within the different FUS condensates. The analysis of the connectivity free energy difference reveals that in both reordered (Fig. 3(e) Left) and arginine-rich mutated (Fig. 3(e) Right) FUS variants, the strength of LARKS–LARKS bonds increases by ∼6 k_*B*_T per amino acid upon ageing (consistent with the atomistic PMF simulations shown in Fig. 1(b), and the increased viscosities from Fig. 3(d)). However, a critical difference between both variants resides on how the location and composition of the strong LARKS-bonds impact the connectivity free energy difference of the rest of amino acids in the sequence. In the reordered FUS variant, the binding energy difference upon ageing for most of the non-LARKS FUS domains is almost negligible (orange colour in 2D projection: ∼0 k_*B*_T, Fig. 3(e) Left). However, in the arginine-rich mutated variant, the abundant formation of tail-located *β*-sheets precludes the interaction among unstructured regions of neighbouring protein replicas, resulting into a severe binding imbalance upon ageing, which in turn, gives rise to a moderately positive free energy differences within non-LARKS domains (red colour in 2D projection: ∼1 k_*B*_T, Fig. 3(e) Right). Such severe imbalance in the inter-molecular forces on top of the unequally distributed strong *vs*. weak patterning (i.e., tail-located LARKS) in the FUS arginine-rich mutated variant and the original sequence (Fig. S7 and Ref.^31^) has been recently shown to drive FUS single-component condensates to display multiphase architectures upon ageing^32,67,111^ or upon phosphorylation^112^. Moreover, this inter-molecular imbalance is consistent with the progressive dynamical arrest of proteins within amorphous condensates observed in LARKS-containing proteins^17,33,36^ such as hnRNPA1^20,64^, FUS^9,67^, TDP-43^90^, or NUP-98^57,91^ among many others^10,20,28,113^.

Furthermore, we apply the primitive path analysis for revealing the transformation of the condensate molecular connectivity across a possible liquid-to-gel transition. Remarkably, even for the reordered FUS variant, which exhibits the lowest viscosity and maturation rate, we detect a notable amount of inter-protein *β*-sheet clusters in the aged condensed phase (Fig. 3(f)). Nevertheless, the isolated *β*-sheet clusters still enable the system to relax as a whole, and behave as a liquid, although with much higher viscosity than pre-aged droplets (Fig. 3(d)). The inter-protein *β*-sheet connectivity of the FUS and mutated FUS sequences (Fig. S3) also display a non-percolated network, although with moderately higher concentration of long-lived *β*-sheet fibrils than the re-ordered FUS variant—consistent with their larger viscosity. Importantly, our results reveal that there is a fundamental requirement for proteins to exhibit complete percolated networks upon maturation: to have at least 2 separate LARKS (with functionality *>*1; i.e., each LARKS can bind to more than one other LARKS: Fig. 2(d)). Nevertheless, their relative location along the sequence is crucial. As shown in Figs. 3(f) and S3, although all the FUS variants studied possess three distinct LARKS, the close location of such LARKS within the low-complexity domain (all contained within a 50-residue length; see Fig. 3(a)) compared to the full-sequence (composed of 526 residues), results in percolation *β*-sheet networks that highly resemble those of the ‘1-Center’ and ‘1-Tail’ low-complexity domain sequences shown in Fig. 2(f) and S2, respectively. This behaviour again highlights the critical importance of LARKS location and abundance in determining the viscoelastic properties of aged condensates, where three equispaced LARKS would likely lead to a complete kinetically trapped network (as recently found for the single low-complexity domain of FUS^11,31^). However, such subtle balance between liquid-like *vs*. partial kinetically arrested behaviour of full-FUS is what possibly gives rise to the recently reported FUS multiphase aged condensates found in vitro^67^ and in silico^32^. Overall, we conclude this Section II D by acknowledging that despite the sequence domain reordering of the low-complexity domain in FUS significantly contributes (by a 25%) to decelerating the rate of inter-peptide *β*-sheet transitions, yet complete ageing inhibition is not fully achieved.

### E. RNA can suppress ageing in domain-reordered FUS condensates

The viscoelastic properties of RNA-binding protein condensates are largely regulated by the presence of RNA in a sophisticated manner^114,115^. RNA concentration, structure, sequence, and chain length deeply control both stability and kinetics of phase-separated condensates^19,20, 62,116–118^. For instance, RNA is known to induce a concentration-dependent reentrant phase behaviour^92,104,^ for a long list of RNA-binding proteins— including, FUS^92,104,105^, PR_25_^47^, G3BP1^63^, LAF-1^119^, or Whi3^120^, among many others^62,121^. Moreover, while short single-stranded disordered RNA can notably reduce condensate viscosity (as observed in LAF-1, an RBP critical in the formation of P granules^119^), long RNAs can also increase viscosity in a concentration-dependent manner^120,122^. Importantly, the LLPS behaviour of numerous RBPs containing low-complexity domains—which exhibit progressive maturation over time and are connected with the onset of several neurodegenerative diseases^25,123^—is widely regulated by RNA. Major examples of these proteins are FUS^18,98,99^, hnRNPA1^7,124^ TAF-15^103,106^, EWSR1^19,103,106^ or TDP-43^125–127^. We focus on FUS since it is an aggregation-prone protein which can undergo LLPS assembling into kinetically trapped biomolecular condensates^25^. Specifically, pathological deposits of FUS-aggregates have been found as hallmarks of diverese neurodegenerative diseases such as FTD and ALS^25,60,73,128^. Since the two previously proposed modifications for the FUS sequence (Fig. 3(a)) are likely to display enhanced RNA-binding affinity through the LARKS adjacent regions, in this section we now investigate the impact of single-stranded RNA in the accumulation of inter-protein *β*-sheets within droplets of the distinct FUS variants.

We begin by studying the RNA-driven reentrant phase behaviour of the three different FUS proteins (Fig. 4(a)). By performing Direct Coexistence simulations of the three different FUS systems in the presence of increasing concentrations of poly-Uridine (polyU) strands of 125 nucleotides (nt), we first determine the maximum temperature (i.e., the critical temperature) at which phase separation occurs. We choose polyU (modelled through the HPS-compatible RNA force field^49^) as it has been largely employed as a model for disordered single-stranded RNA both *in vitro*^103,105,119,129^ and *in silico*^31,85,117^ studies of RNA-binding protein condensates. The critical temperature (inversely related to the protein saturation concentration^85^) is plotted as function of the polyU/FUS mass ratio for the three FUS sequences in Fig. 4(a). While at very low RNA concentration, the stability of the condensates moderately increases, at higher RNA concentrations the phase-separated droplets are destabilized— in agreement with experimental findings^62,92,104,105^. We note that the length of the RNA can shift significantly the concentration thresholds at which the different behaviours in Fig. 4(a) emerge^119,130^. However, for FUS– RNA condensates, the impact of RNA length is negligible up to high RNA concentrations^117^, and phase separation remains stable as a function of length, as long as the RNA chains are of at least 125-nt in length^85^—hence, our choice of 125-nt strands. Interestingly, we find that the two proposed FUS variants can enhance condensate stability up to moderately higher RNA concentrations than the unmodified FUS protein (e.g., ∼25% higher polyU/FUS mass ratio than the original sequence), which is indicative of their higher RNA-binding affinity. For the arginine-rich mutated variant, this is an expected result since the number of arginines is higher than in the original sequence (Fig. 3(a)). However, for the domain-reordered variant, that is a remarkable finding, given that the amino-acid composition of the protein has been kept unaltered. Such sensitivity of FUS phase behaviour to the location of the LCD again highlights the critical importance not only of the sequence composition of a given binding motif, but also of the sequence context in which a given motif is found. Furthermore, the condensates of the relocated FUS variant not only incorporate higher polyU concentrations while maintaining their stability, but also exhibit a higher enhancement of condensate stability at low RNA concentration than the original FUS sequence and its arginine-rich mutated version (Fig. 4(a)).

**Figure 4:**
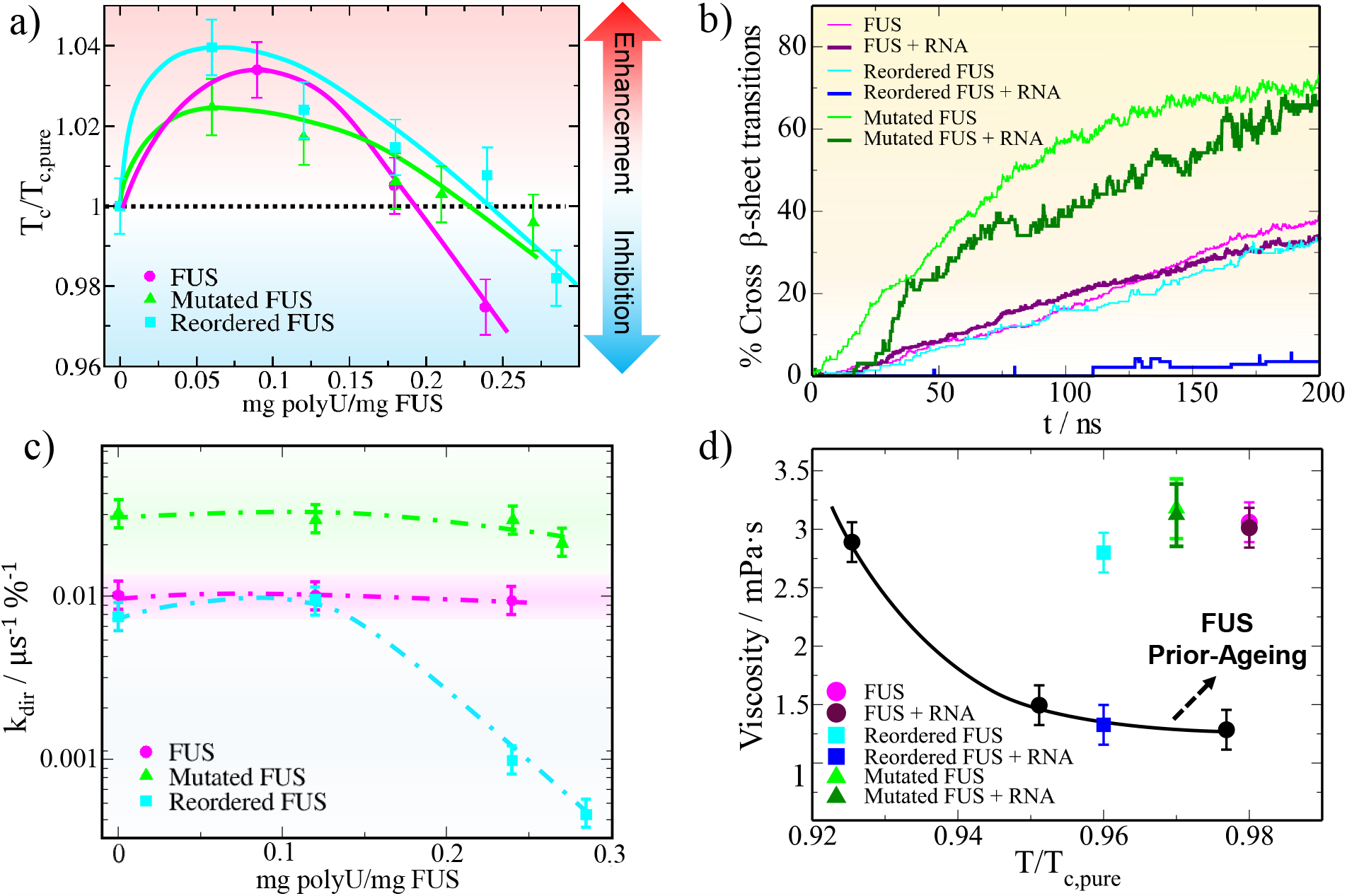
RNA at moderate concentration inhibits the formation of inter-protein *β*-sheet clusters in sequence domain-reordered FUS condensates. a) Critical temperature (filled symbols) of FUS (magenta), mutated FUS (green) and reordered FUS (cyan) condensates as a function of the polyU/protein mass ratio. Symbols above the horizontal dotted line indicate RNA-driven LLPS enhancement while symbols below such threshold pinpoint inhibition of condensate phase-separation. Continuous curves are included as a guide for the eye. Please note that temperatures have been renormalized by the critical temperature of each sequence in absence of RNA (*T*_*c,pure*_). b) Time-evolution of the percentage of LARKS forming crossed-*β*-sheet motifs in phase-separated condensates of the three FUS variants both in absence and presence of RNA as indicated in the legend. The polyU/protein mass ratio of each condensate corresponds to 0.24, 0.27, and 0.28 for FUS, mutated FUS and reordered FUS, respectively (i.e., that is the maximum polyU concentration at which condensates are stable above *T/T*_*c,pure*_ ∼0.97; temperature at which all simulations were performed). c) Estimated kinetic constants (from a second-order reaction analysis to the number of inter-protein *β*-sheet transitions over time) for the different polyU/FUS condensates at increasing polyU/protein mass ratios at *T* ∼0.97 *T*_*c,pure*_. Error bars account for the estimated uncertainty while dotted lines are included as a guide for the eye. d) Viscosity as a function of temperature for pure FUS condensates before the emergence of inter-protein *β*-sheet transitions (black circles), and after a maturation time interval of 0.3 *μ*s for each FUS variant both in absence and presence of RNA as indicated in the legend. Temperature has been renormalized by *T*_*c,pure*_ of each FUS variant. The polyU/protein mass ratios of the mixed condensates are the same as those described in Panel (b).

We now ask how RNA might influence the progressive formation of inter-protein *β*-sheet motifs in phase-separated droplets. To the different solutions of FUS proteins, we now add the maximum concentration of RNA that still enables phase separation near the critical temperature of the FUS condensates without RNA (i.e., *T/T*_*c,pure*_ *>* 0.95). Then, in bulk conditions and at the coexistence droplet density of each mixture at *T/T*_*c,pure*_ ∼ 0.97, we measure the number of structural cross-*β*-sheet transitions over time (Fig. 4(b)). Remarkably, whereas the accumulation of inter-protein *β*-sheets for the mutated FUS sequence is only moderately lower than in absence of polyU, for the reordered FUS variant, RNA inhibits the emergence of LARKS disorder-to-order structural transitions within the condensates (Fig. 4(b); dark blue curve). For the original FUS sequence, polyU only slightly decelerates the gradual formation of inter-protein *β*-sheets at moderate RNA concentration^31^ (i.e., as in the mutated variant). These results strongly suggest that, besides LARKS location within the sequence, the chemical make-up of the neighboring domains, which in FUS reordered display high binding affinity for RNA, are critical to control LARKS–LARKS high-density fluctuations needed to trigger inter-protein *β*-sheet transitions. The combination of RNA binding to the regions adjacent to the low complexity domain (i.e., RRMs or RGGs)—which severely reduces the likelihood of inter-protein interactions—and RNA–RNA long-range electrostatic repulsion in the vicinity of the FUS low-complexity domain, prevents the spontaneous formation of high-density LARKS clusters. Nevertheless, given that in the original and mutated FUS variants, the RRM and RGG domains are all positioned towards the mid/end of the sequence (Fig. 3(a)), this effect of RNA in preventing the spontaneous formation of low-complexity domain high-density clusters is much milder than in the reordered variant. According to the fast formation of *β*-sheet clusters in the mutated sequence, we can conclude that RNA is not establishing strong contacts with the positively charged mutated patches near the three consecutive LARKS in the FUS low-complexity domain (Fig. 5(a)). Possibly, the binding affinity of RNA for the RRM and RRG domains is still much stronger than that for the arginine mutated regions within the low-complexity domain^115,131^. Therefore, RNA exhibits a weak effect in blocking low-complexity domain density fluctuations in the arginine-rich mutant. The effects observed in the arginine-rich mutant might have been accentuated by the overestimation of cation-*π* interactions, respect to electrostatics interactions, in the employed force field^85,101^. Such imbalance in the force field could contribute to undermining the effectivity of the mutated FUS sequence in avoiding low-complexity domain high-density fluctuations through self-repulsive R–R interactions and favourable R–U interactions in presence of RNA.

**Figure 5:**
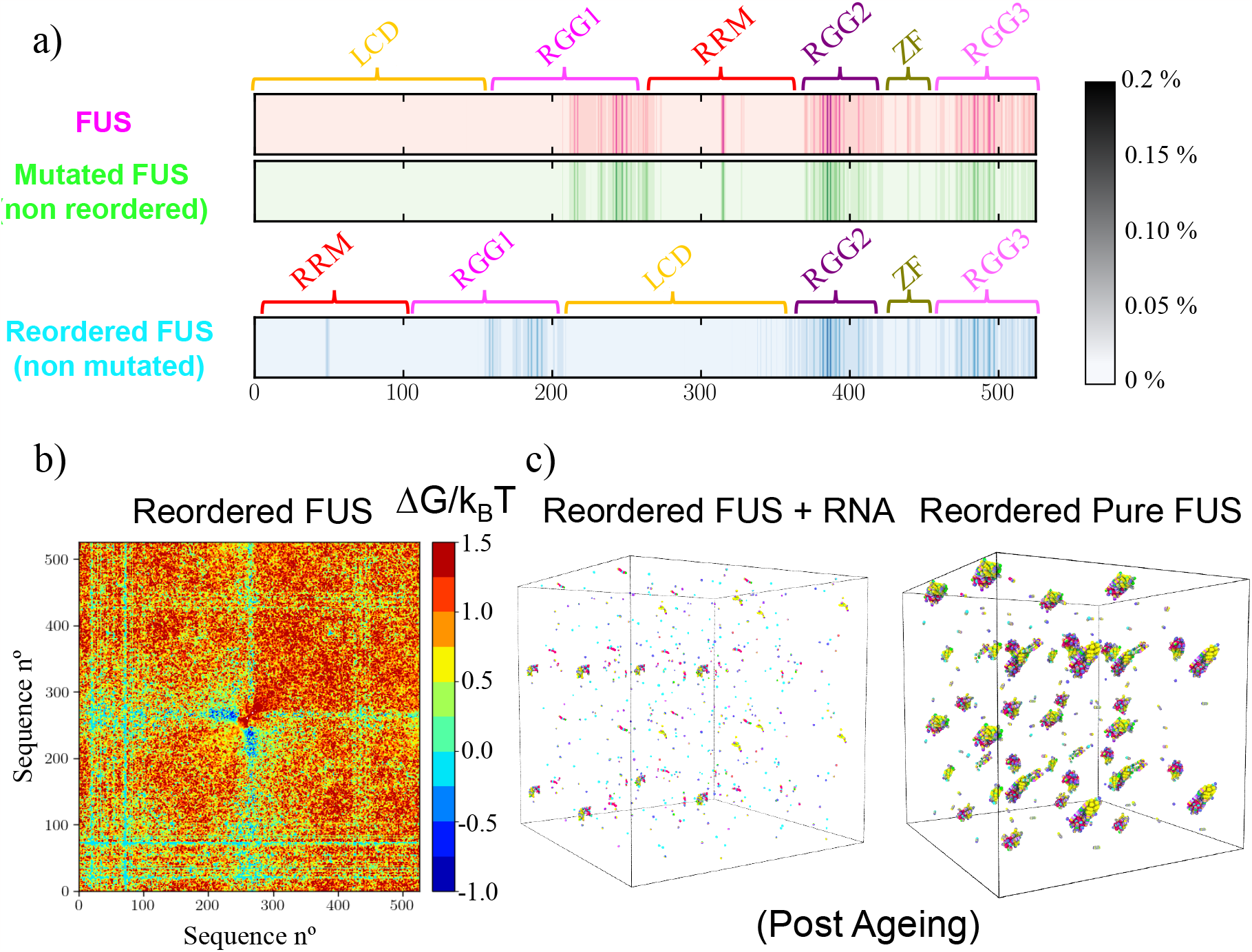
The location of the strong RNA-binding domains in FUS is compelling to inhibit disorder-to-*β*-sheet transitions at moderate RNA concentrations. a) Average contact probability per residue (scale bar) between polyU and the different amino acids composing the three FUS variants in phase-separated condensates pre-ageing at *T/T*_*c,pure*_ ∼0.97 and the maximum polyU/protein mass ratio at which condensates are stable at such temperature (i.e., 0.24 for FUS, 0.27 for mutated FUS, and 0.28 for the reordered FUS variant). b) Free energy inter-molecular variation computed from the molecular contact probability between aged reordered FUS pure condensates and polyU/reordered FUS ones at a polyU/protein mass ratio of 0.28, *T* ∼0.97 *T*_*c,pure*_, and after an ageing time interval of 0.3 *μ*s for both systems. c) *β*-sheet network connectivity (evaluated through the PPA method) of an aged polyU/reordered FUS condensate (Left) and a pure reordered FUS condensate (Right) upon 0.3 *μ*s of maturation.

To better comprehend the effect of RNA insertion in FUS droplet maturation, we evaluate the kinetic constants of *β*-sheet formation for the three FUS sequences as a function of the polyU/FUS mass ratio (Fig. 4(c)). For all sequences, *k*_*dir*_ roughly varies while adding low concentrations of polyU compared to the pure condensate. However, at higher, although still moderate, RNA concentration (0.28 polyU/FUS mass ratio), the structural transition kinetic constants of the reordered FUS variant drop by more than one order of magnitude. On the contrary, for the mutated and original sequences *k*_*dir*_ is only moderately reduced by ∼30% and ∼10%, respectively, at the maximum RNA concentration that condensates admit without being destabilised. This RNA concentration-dependent behaviour found for all sequences (especially for the reordered variant) highlights the key role of long-range electrostatic repulsion of RNA– RNA in reducing droplet density and thus preventing high-density protein fluctuations (which can be only attained at moderate RNA concentration)^31^. Moreover, these results pinpoint the major role of RNA-binding to LARKS neighbouring domains.

If we now evaluate the viscosity of aged condensates in both the absence *vs*. presence of RNA after the same maturation time (i.e., 0.3 *μs*), we observe that the increase in viscosity for the original and mutated FUS proteins is barely the same with or without polyU (Fig. 4(d); please note that the same RNA concentrations as those employed in Fig. 4(b) have been used here). However, for the FUS reordered variant, the viscosity is ∼2.5 times lower in the presence of RNA than for pure aged condensates. Impressively, such value (dark blue square) is very similar to those obtained for pure FUS condensates prior to ageing at the same temperature. The viscosity dependence with temperature of FUS droplets prior to ageing is remarkably similar to that of mutated and reordered pre-aged condensates (black curve in Fig. 4(d)). Whereas the reordering of the LCD in pure FUS droplets just led to a moderate deceleration in the structural transition rate (Fig. 3(c)) and to a slight decrease in viscosity compared to the mutated and non-modified FUS sequences, in the presence of RNA, the accumulation of *β*-sheet content over time is strongly inhibited (Fig. 4(b)). For mixtures of the different FUS sequences, both in the absence or presence of RNA, we would expect an intermediate ageing behaviour with respect to the pure variant systems.

To characterise from a molecular perspective the crucial role of RNA in suppressing ageing of LCD-reordered FUS condensates, we evaluate the binding probability of polyU with FUS for pre-aged condensates of each variant studied (Fig. 5(a)). We find that for all FUS-RNA systems, the regions that mostly interact with polyU are the three RGG domains followed by the RNA-recognition motif and the zinc finger. In contrast, the low-complexity domain barely establishes interactions with polyU in all the FUS proteins studied. Such a negligible binding probability of the low-complexity domain to RNA would demand that at least its neighbouring domains could establish substantial protein–RNA interactions upon RNA addition, to hinder the accumulation of *β*-sheets. Although that is the case for the reordered FUS variant, in both FUS and its mutated version, there is a gap of approximately 140 residues from the three consecutive LARKS in the low-complexity domain and the first region of the RGG1 domain that exhibits substantial interactions with RNA. Furthermore, the three LARKS are positioned near one of the protein terminal regions, which also contributes to speeding up the formation of *β*-sheets over time (as shown in Fig. 2(b) for the ‘1-Tail’ sequence). Oppositely, in the reordered FUS variant, the central location of the LCD—also flanked by two highly RNA-interacting RGG domains—contributes to impeding the spontaneous fluctuations needed to bring multiple LARKS close. For the reordered sequence, we also determine the contact probability binding ratio (see Supplementary Material for further details on this calculation) between aged condensates in the absence *vs*. presence of RNA (at moderate concentration) after the same maturation timescale (0.3 *μs*). As shown in Fig. 5(b), the net variation in protein binding probability upon RNA inclusion is positive, and the binding free energy increases by ∼1.5 k_*B*_T on average for most of the sequence regions (implying lower protein binding probability). Importantly, such behaviour emerges despite the RNA not being able to directly target the LCD, and instead binding the adjacent arginine-glycine rich-regions (Fig. 5(a)). The only regions that exhibit an increase in their binding probability with respect to aged droplets in the absence of RNA are those that can now interact with the unstructured LARKS domains released from inter-protein *β*-sheets. Yet, the overall free energy increase collectively hinders local high-density fluctuations within FUS-RNA condensates that underlie the progressive accumulation of inter-protein *β*-sheet motifs^14,37,38,56^. Taken together, these results highlight that the specific binding affinity between the adjacent LCD domains and RNA contribute to frustrating the formation of inter-protein *β*-sheets. Thus, the addition of polyU in combination with a sequence domain reordering in which the LCD-flaking domains do not exhibit substantial RNA-binding affinity might not effectively preclude inter-protein structural transitions.

Finally, we compare the *β*-sheet percolation network of aged reordered FUS condensates both in absence and presence of polyU (Fig. 5(c)), to explain the remarkably similar viscosities of aged droplets with moderate RNA concentration and pure droplets pre-ageing (Fig. 4(d)). As quantified in Fig. 4(c) and illustrated in Fig. 5(c), RNA inclusion in reordered FUS droplets reduces the kinetic constant of structural transitions by more than one order of magnitude, and suppresses the emergence of highly dense *β*-sheet networks that induce extremely low protein diffusivity. Taken together, these results reveal how the simple reordering of a protein sequence, which may not even initially affect its ability to phase-separate, can have a tremendous impact on the long-term viscoelastic properties of the condensate, and therefore, in driving it from functional to pathological. Furthermore, we also demonstrate here how RNA can act as a potent regulator of LLPS^19,20,62,85,116,117^ by specifically binding to protein regions that largely control, directly or indirectly, the macroscopic phase behaviour of biomolecular condensates.

## III. CONCLUSION

In this work, we have developed a multiscale computational approach—combining atomistic simulations and coarse-grained models of different resolutions—to elucidate the impact of the abundance and location of LARKS in ageing of biomolecular condensates. The aim of our simulations is to predict effective strategies to inhibit LARKS inter-protein structural transitions precluding liquid-like behaviour in RBP condensates^14,28,31,56^. Based on structured *vs*. disordered peptide binding energies predicted from all-atom PMF calculations (Fig. 1(a-b)), we first develop a coarse-grained model for LARKS-containing low-complexity domains that progressively undergo *β*-sheet fibrillization in phase-separated condensates (Fig. 1(d)). Through this model, we reveal how the abundance and location of LARKS imperatively determine the maturation rate, defined as the number of structural disorder-to-*β*-sheet transitions over time. We find that low-complexity domain sequences with LARKS located near the extremes of the protein exhibit much faster maturation rates than those in which LARKS are placed towards the center (Fig. 2(b)). Remarkably, low-complexity domains with a tail-located LARKS displayed much higher viscosity after a given window of time than those in which the LARKS is placed towards the center. Yet, at very long timescales, low-complexity domains with a single LARKS (independently of its location) can still relax and behave as a high viscous liquid. However, phase-separated condensates of proteins containing two or more LARKS become kinetically trapped and display gel-like behaviour upon maturation (Figs. 2(c-d)).

Condensate rigidification is also correlated with moderate densification over time, as pinpointed by our results, and recently reported by *in silico*^53^ and *in vitro* studies^11^. Furthermore, our simulations reveal that the accumulation of inter-peptide *β*-sheets, which can gradually transform weak and transient network connectivities into almost irreversible long-lived percolated networks (Fig. 2(f)), strongly depends on both abundance and LARKS location. Hence, identifying different sequence patterns that present low maturation rates and allow protein condensates to maintain functional liquid-like behaviour for longer timescales might be a potential framework to design strategies to decelerate aberrant phase transitions driven by *β*-sheet formation^25,60^.

Furthermore, we show how a simple reordering of the different FUS domains can decelerate the maturation rate of pure condensates by a 30% without altering its ability to phase-separate (Fig. 3). Such reordering is effective because the 3-LARKS-containing region is placed in the center of the protein, and is flanked by the three native RGG domains of FUS. In contrast, S → R and G → R mutations of the adjacent FUS LARKS residues, intended to disrupt LARKS *β*-sheet binding, substantially enhance the formation of structural transitions over time. Such increase in the *β*-sheet rate directly translates into higher condensate viscosities than those of the original sequence and the reordered FUS variant after similar maturation timescales (Fig. 3(d)). However, when adding moderate concentrations of RNA (polyU) to the reordered FUS sequence, we strikingly find that the formation of inter-protein *β*-sheet content is now inhibited (Figs. 4 and 5). The molecular mechanism behind such extreme deceleration in inter-protein *β*-sheet formation is an RNA-blocking mechanism, which impedes the spontaneous formation of high-density clusters of LARKS (Fig. 5). As a consequence, aged condensates of the FUS re-ordered sequence in presence of RNA exhibit similar viscosities to those of FUS prior-ageing. Contrarily, the original and mutated FUS sequences with RNA, upon the same maturation timescale increased their viscosity by 300% compared to pre-aged FUS condensates (Fig. 4(d)). Taken together, our work uncovers, from a molecular perspective the role of LARKS abundance and location in facilitating or hindering ageing of RNA-binding proteins^9,10,19,20,91^.

## ACKNOWLEDGMENTS

This project has received funding from the Oppenheimer Research Fellowship of the University of Cambridge. S. B. acknowledges funding from the Spanish Ministry of Economy and Competitivity (PID2019-105898GB-C21) and the Oppenheimer Fellowship. A. T. is funded by the Universidad Politécnica de Madrid through the PhD fellowship (‘programa propio UPM’) and the Oppenheimer Fellowship. I. S.-B. acknowledges funding from the Oppenheimer Fellowship, Derek Brewer scholarship of Emmanuel College and EPSRC Doctoral Training Programme studentship, number EP/T517847/1. J. R. and M. M. C. acknowledge funding from the Spanish Ministry of Economy and Competitivity (PID2019-105898GA-C22) and the Madrid Government (Comunidad de Madrid-Spain) under the Multiannual Agreement with Universidad Politécnica de Madrid in the line Excellence Programme for University Professors, in the context of the V PRICIT (Regional Programme of Research and Technological Innovation). M.M.C. also acknowledges CAM and UPM for financial support of this work through the CavItieS project No. APOYO-JOVENES-01HQ1S-129-B5E4MM. J. R. E. also acknowledges funding from the Roger Ekins Research Fellowship of Emmanuel College, and the Ramon y Cajal fellowship (RYC2021-030937-I). R.C.-G. acknowledges funding from the European Research Council (ERC) under the European Union Horizon 2020 research and innovation programme (grant agreement 803326). This work has been performed using resources provided by the Cambridge Tier-2 system operated by the University of Cambridge Research Computing Service (http://www.hpc.cam.ac.uk) funded by EPSRC Tier-2 capital grant EP/P020259/1. The authors gratefully acknowledge the Universidad Politécnica de Madrid (www.upm.es) for also providing computing resources on Magerit Supercomputer.

## DATA AVAILABILITY

The data that supports the findings of this study are available within the article and its Supplementary Material. The LAMMPS and GROMACS files of the residue-resolution models and all-atom simulations respectively, as well as the dynamic algorithm software are available upon reasonable request to the corresponding authors of the article.

## COMPETING INTERESTS

The authors declare no competing interests.

## Notes

### Competing Interest Statement

The authors have declared no competing interest.

### Summary of Updates

This version of the manuscript has been revised to update the comments adressed by the reviewers of this work.

