## Supporting Information for "Location and concentration of aromatic-rich segments dictates the percolating inter-molecular network and viscoelastic properties of ageing condensates"

Samuel Blazquez[1,2], Ignacio Sanchez-Burgos[2], Jorge Ramirez[3], Tim Higginbotham[2], Maria M. Conde[3], Rosana Collepardo-Guevara[2,4,5,*], Andres R. Tejedor[2,3,*] and Jorge R. Espinosa[1,2,*]
*[1] Department of Physical-Chemistry, Universidad Complutense de Madrid, Av. Complutense s/n, 28040, Madrid, Spain*
*[2] Maxwell Centre, Cavendish Laboratory, Department of Physics, University of Cambridge, J J Thomson Avenue, Cambridge CB3 0HE, United Kingdom.*
*[3] Department of Chemical Engineering, Universidad Politécnica de Madrid, José Gutiérrez Abascal 2, 28006, Madrid, Spain.*
*[4] Yusuf Hamied Department of Chemistry, University of Cambridge, Lensfield Road, Cambridge CB2 1EW, United Kingdom*
*[5] Department of Genetics, University of Cambridge, Cambridge, CB2 3EH*
** = To whom correspondence should be sent.*
(Dated: June 9, 2023)

### SI. MODELS AND SIMULATION DETAILS

#### A. Atomistic simulations for evaluating potential of mean force binding energies

We estimate the potential of mean force (PMF) by performing a set of atomistic Umbrella Sampling Molecular Dynamics (MD) simulations for the NFGAFS segment (PDB code: 5WHN) of the TDP-43 low-complexity domain (LCD). All-atom PMF simulations are performed with explicit water and ions using the a99SB-*disp* force field for proteins, water and ions [1] and the GROMACS 2018 MD package [2]. The starting configuration for each simulation consisted of six stacked peptides (of the same sequence; three pairs of peptides stacked on top of each other forming a ladder) taken from the cross-$\beta$-fibril structure resolved crystallographycally in Ref. [3]. Acetyl and N-methyl capping groups were added to the termini of each peptide. For each 6-peptide system, we perform two sets of simulations. First, we compute the dissociation binding energy of a single structured peptide from a 6-peptide aggregate when it exhibits the cross-$\beta$-sheet structure (imposed through positional restraints on all the heavy atoms of 1000 kJ mol$^{-1}$ nm$^{-2}$ in all directions, except for the dissociating peptide in the pulling direction; solid green curve in Figure 1(a) of the main text). This ensures that the six individual peptides maintain their secondary structure along the dissociation pathway. Then, in the second set of simulations, we compute the dissociation binding PMF profile among disordered peptides, *i.e.*, by enabling all peptides to freely sample their configurational landscape (dashed red curve in Figure 1(a) of the main text). Importantly, in both sets of simulations, we kept the position of the central $C_\alpha$ atom of each peptide fixed to control the relative distances between peptides (except in the direction of the reaction coordinate for the dissociating peptide).

For the reaction coordinate, we use the distance between the center of mass (COM) of the 'dissociating' peptide and the closest peptide to it along the pulling direction. In the case of non-structured peptides, rather than the COM distance, we employ the distance between the central $C_\alpha$'s of the aforementioned chains to avoid noise due to changes in the COM from fluctuations in the conformation of the peptide. We space umbrella windows approximately every 0.2 Å along the reaction coordinate, from 4 to 30 Å (where long-range interactions (*i.e.*, electrostatic interactions) completely vanish). To sample the steep potential among rigid structures, a spring constant of 24000 kJ mol$^{-1}$ nm$^{-2}$ is used along the entire dissociation pathway. For consistency, the same value of the spring constant is also employed in the disordered PMF calculations.

For the structured peptides, we run simulations of about 30 ns per umbrella window. For disordered peptides, a simulation timescale of 100 ns is necessary. Such timescale guarantees uncertainties of about 1.5 and 3 $k_B$T at most in the dissociation profiles of the disordered and structured LARKS respectively. For the integration of the equations of motion, we employ the Verlet algorithm with a time step of 1 fs. Bond lengths are constrained by using the LINCS algorithm [4] with an order of 8 and 2 iterations. The cut-off for the Coulomb and van der Waals interactions was fixed at a value of 14 Å. For electrostatics, we use Particle Mesh Ewald (PME) [5] of fourth order with a Fourier spacing of 0.12 nm and an Ewald tolerance of $1.5{\cdot}10^{-5}$. Simulations were performed in the NpT ensemble. To keep

| | A | | | | | B | | | | | C | | | | | D | | | | |
|---|---|---|---|---|---|---|---|---|---|---|---|---|---|---|---|---|---|---|---|---|
| | $\sigma$/Å | $\epsilon$/kcal·mol$^{-1}$ | $\nu$ | $\mu$ | r$_c$/Å | $\sigma$/Å | $\epsilon$/kcal·mol$^{-1}$ | $\nu$ | $\mu$ | r$_c$/Å | $\sigma$/Å | $\epsilon$/kcal·mol$^{-1}$ | $\nu$ | $\mu$ | r$_c$/Å | $\sigma$/Å | $\epsilon$/kcal·mol$^{-1}$ | $\nu$ | $\mu$ | r$_c$/Å |
| A | 5.88 | 0.255 | 1 | 1 | 11.8 | 5.88 | 0.255 | 1 | 1 | 11.8 | 5.88 | 0.255 | 1 | 1 | 11.8 | 5.88 | 0.255 | 1 | 1 | 11.8 |
| B | | | | | | 5.25 | 1.53 | 1 | 3 | 21.0 | 5.88 | 0.255 | 1 | 1 | 11.8 | 5.88 | 0.255 | 1 | 1 | 11.8 |
| C | | | | | | | | | | | 5.25 | 1.53 | 1 | 3 | 21.0 | 5.88 | 0.255 | 1 | 1 | 11.8 |
| D | | | | | | | | | | | | | | | | 5.25 | 1.53 | 1 | 3 | 21.0 |

**TABLE S1:** Model parameters for the different $\sigma$, $\epsilon$, $\mu$, $\nu$ and $r_c$ values of our LCD sequences shown in Figure 1(c) of the main text. In all cases, the mass of every bead/residue is 118.9 g/mol.

constant temperature and pressure we use the Nosé–Hoover thermostat [6] at T = 300 K (with 1 ps relaxation time) and the Parrinello–Rahman [7] isotropic barostat at p = 1 bar (with a 1 ps relaxation time), respectively. A NaCl concentration of 0.15 M was used throughout all our simulations.

Each simulation system comprises a volume size of approximately 7 x 8 x 6 nm$^3$. We simulate about 10000 water molecules. All of the systems used were electroneutral (*i.e.*, the total net charge was zero). After solvation of the peptides, we perform an energy minimization with a force tolerance of 1000 kJ mol$^{-1}$ nm$^{-1}$, followed by a short, 1000 ps NpT equilibration both with positional restraints (of 10000 kJ mol$^{-1}$ nm$^{-2}$) for the heavy atoms of the chains in all three directions of space, except for the capping groups, which were allowed to rotate and freely relax into the lowest potential energy configuration. We analyze the Umbrella Sampling simulations using the WHAM [8] tool implemented in GROMACS. The first 10% of the simulations were discarded as the equilibration time.

#### B. LCD coarse-grained simulations

We develop an innovative coarse-grained model for generic LCD sequences in which we integrate structured vs. disordered peptide binding energies from our atomistic simulations (Figure 1(a) and 1(b) of the main text). We model the different studied low-complexity domains with a total sequence length of 125 residues (which is the typical length of a low-complexity domain in a phase-separating RNA-binding protein (RBP) [9, 10]). To describe non-bonded interactions between the different LCD residues, we employ the Wang-Frenkel (WF) potential which is described by the following equation:

$$\phi(r) = \epsilon\alpha \left[ \left(\frac{\sigma}{r}\right)^{2\mu} - 1 \right] \left[ \left(\frac{r_c}{r}\right)^{2\mu} - 1 \right]^{2\nu},$$

with

$$\alpha = 2\nu \left(\frac{r_c}{\sigma}\right)^{2\mu} \left[ \frac{1+2\nu}{2\nu\left[(r_c/\sigma)^{2\mu}-1\right]} \right]^{2\nu+1},$$

and

$$r_{min} = r_c \left[ \frac{1+2\nu}{1+2\nu(r_c/\sigma)^{2\nu}} \right]^{1/2\nu},$$

where $\epsilon$ is the depth of the WF potential, $r$ represents the distance between two amino acids, and $\sigma$ accounts for the molecular diameter of each amino acid. $r_c$ is the cut-off distance of the potential, and $\alpha$ is a coefficient that ensures that the depth of the attractive well is $\epsilon$. $\nu$ and $\mu$ are positive integers. The mass of each bead was chosen to be m = 118.9 g/mol. The parameters ($\sigma$, $\epsilon$, $\nu$, $\mu$ and $r_c$) for the different residue-residue interactions of our LCD simulations are shown in Table S1.

Bonds between consecutive beads are modelled with a harmonic potential $V_{\rm har} = K(r - r_0)^2$ of $K = 23.87$ kcal/(mol·Å$^2$), and $r_0$ of 5.88 Å. Furthermore, we apply an angular potential of the form, $V_{\rm ang} = K_\theta\ (\theta–\theta_0)^2$, between consecutive bonds, with an angular constant of $K_\theta = 1 \cdot 10^{-5}$ kcal/mol and a resting angle of $\theta_0 = 180°$ for fully disordered chain replicas, and with a constant of $K_\theta = 5$ kcal/mol and $\theta_0 = 180°$ for structured LARKS residues when the inter-peptide $\beta$-sheet transition has taken place. The values of $\epsilon$ and $\mu$ after the transition are chosen to reproduce the increase in the binding energy between the structured domains in the PMFs (see Fig. S1 in [11] for a similar mapping from disordered to order interactions).

System sizes in our LCD coarse-grained simulations contained 81 replicas of each low-complexity domain sequence. We perform Direct Coexistence simulations [12] to compute the phase diagram of the LCDs in the NVT ensemble (*i.e.*, constant number of particles (N), volume (V) and temperature (T)). For all of our simulations, we have used the LAMMPS Molecular Dynamics package [13]. The Nosé–Hoover thermostat [14–16] with a relaxation time of 0.5 ps has been applied to keep the temperature constant. We have also applied periodic boundary conditions in the three directions of space. Verlet integration of the equations of motion has been used with an integration timestep of 5 fs. We have studied the disorder-to-order transitions of the different LCD sequences through our dynamical ageing model (further details in Section SIII) in bulk phases running NpT simulations of cubic boxes using the Nosé-Hover thermostat [6] and Parrinello–Rahman [7] barostat at $T/T_c = 0.91$ and $p = 0$ bar (both with a relaxation time of 0.5 ps). The viscoelastic properties have been evaluated through cubic NVT simulations. The volume of each box is set to the bulk equilibrium density of the condensate at the corresponding temperature. The typical length of each simulation required to compute the viscosity is about 3 $\mu$s.

The sequences proposed in this work (Fig. 1(c) of the main text) to study the effect of LARKS abundance and location in protein phase separation are the following:

**1-Tail**

AAAABBBBBBAAAAAAAAAAAAAAAAAAAAAAAAAAAAAAAAAAAAAAAAAAAAAAAAAAAAAAAAAAAAAAAAAAAAAA
AAAAAAAAAAAAAAAAAAAAAAAAAAAAAAAAAAAAAAAAAAAAA

**1-Center**

AAAAAAAAAAAAAAAAAAAAAAAAAAAAAAAAAAAAAAAAAAAAAAAAAAAAAAAAAAABBBBBBAAAAAAAAAAAAAAA
AAAAAAAAAAAAAAAAAAAAAAAAAAAAAAAAAAAAAAAAAAAAA

**2-OppositeTail**

AAAABBBBBBAAAAAAAAAAAAAAAAAAAAAAAAAAAAAAAAAAAAAAAAAAAAAAAAAAAAAAAAAAAAAAAAAAAAAA
AAAAAAAAAAAAAAAAAAAAAAAAAAAAAAAAAAACCCCCCAAAA

**2-SameTail**

AAAABBBBBBAAAAAAAAAAAAAACCCCCCAAAAAAAAAAAAAAAAAAAAAAAAAAAAAAAAAAAAAAAAAAAAAAAAAA
AAAAAAAAAAAAAAAAAAAAAAAAAAAAAAAAAAAAAAAAAAAAA

**2-Center**

AAAAAAAAAAAAAAAAAAAAAAAAAAAAAAAAAAAAAAAAAAAAAAAAABBBBBBAAAAAAAAAAAAAACCCCCCAAAAA
AAAAAAAAAAAAAAAAAAAAAAAAAAAAAAAAAAAAAAAAAAAAA

**2-Equispaced**

AAAAAAAAAAAAAAAAAAAAAAAAAAAAAAAAAAAAABBBBBBAAAAAAAAAAAAAAAAAAAAAAAAAAAAAAAAAAAAA
CCCCCCAAAAAAAAAAAAAAAAAAAAAAAAAAAAAAAAAAAAAAA

**3-Equispaced**

AAAAAAAAAAAAAAAAAAAAAAAAAABBBBBBAAAAAAAAAAAAAAAAAAAAAAAAAACCCCCCAAAAAAAAAAAAAAAA
AAAAAAAAAADDDDDDAAAAAAAAAAAAAAAAAAAAAAAAAAAAA

**3-SameTail**

AAAABBBBBBAAAAAAAAAAAAAACCCCCCAAAAAAAAAAAAADDDDDDAAAAAAAAAAAAAAAAAAAAAAAAAAAAAAA
AAAAAAAAAAAAAAAAAAAAAAAAAAAAAAAAAAAAAAAAAAAAA

where A represents a generic non-LARKS unstructured residue, and B, C, and D are LARKS residues capable of undergoing a structural $\beta$-sheet transition. Please note that only inter-protein $\beta$-sheet aggregation can occur between residues of the same type (*i.e.*, B-B, C-C, or D-D).

#### C. Residue-resolution coarse-grained simulations with the HPS-Cation-$\pi$ model

In order to simulate the different studied variations of the FUS sequence, we employ the recent reparameterization by Das *et al.* [17] of the chemically-accurate coarse-grained (CG) HPS protein model proposed by Dignon *et al.* [18] implemented in the LAMMPS Molecular Dynamics package [13]. For RNA chains, we use the HPS-compatible CG model proposed by Regy *et al.* [19]. The coarse-grained model resolution, both for FUS and RNA, is of one bead per amino acid and nucleotide respectively. In this model, the intrinsically disordered regions (IDRs) of the proteins are considered as fully flexible polymers, while the structured globular domains are treated as rigid bodies (where their conformations are taken from the Protein Data Bank (PDB) crystalline structure – see SI D for the PBD codes) by using the rigid body integrator of LAMMPS [13]. Besides, structured globular domains interactions are scaled down by a 30% to account for the 'buried' amino acids as proposed by Krainer *et al.* [20]. RNA strands are also treated as flexible polymers. The potential energy of proteins and nucleic acids modelled through the HPS-Cation-$\pi$ force field is given by:

$$E = E_{\text{Bonds}} + E_{\text{Electrostatic}} + E_{\text{Hydrophobic}} + E_{\text{Cation}-\pi}, \tag{S1}$$

where $E_{\text{Hydrophobic}}$, $E_{\text{Cation}-\pi}$ and $E_{\text{Electrostatic}}$ interactions are only applied between non-bonded beads and $E_{\text{Bonds}}$ between subsequent bonded beads.

Bonded interactions between adjacent amino acid beads or consecutive RNA nucleotides are described by a harmonic potential:

$$E_{\text{Bonds}} = \sum_{\text{Protein/RNA bonds}} k(r - r_0)^2, \tag{S2}$$

where the equilibrium bond length is $r_0 = 5.0$ Å between subsequent nucleotides and $r_0 = 3.81$ Å between the bonded amino acid beads. The spring constant is $k = 10$ kJ/(mol Å$^2$). Electrostatic interactions, $E_{\text{Electrostatic}}$, between charged amino acids and RNA nucleotides are described by a Yukawa/Debye-Hückel potential of the form:

$$E_{\text{Electrostatic}} = \sum_i \sum_{j<i} \frac{1}{4\pi D} \frac{q_i q_j}{r} e^{-r/\kappa}, \tag{S3}$$

where $D = 80\epsilon_0$ is the dielectric constant of water (being $\epsilon_0$ the vacuum permittivity), $q_i$ and $q_j$ represent the charges of beads $i$ and $j$ (amino acids or nucleotides), $r$ is the distance between the $i$th and $j$th beads, and $\kappa$= 1 nm is the Debye screening length that mimics the implicit solvent (water and ions) at physiological salt concentration ($\sim$150 mM of NaCl) [18].

Hydrophobic interactions between different amino acid types and nucleotides are built upon a scale of amino acid and RNA nucleotide hydrophobicity based on a statistical potential derivation from contacts in PDB structures, and implemented through the functional form of an Ashbaugh/Hatch potential (for further details, see References [18, 19, 21, 22]):

$$E_{\text{Hydrophobic}} = \sum_i \sum_{j<i} \begin{cases} 4\epsilon_{ij} \left[ \left(\frac{\sigma_{ij}}{r}\right)^{12} - \left(\frac{\sigma_{ij}}{r}\right)^{6} \right] + (1-\lambda_{ij})\epsilon_{ij}, & r < 2^{1/6}\sigma_{ij} \\ \lambda_{ij} 4\epsilon_{ij} \left[ \left(\frac{\sigma_{ij}}{r}\right)^{12} - \left(\frac{\sigma_{ij}}{r}\right)^{6} \right], & \text{otherwise}, \end{cases} \tag{S4}$$

where $\lambda_i$ and $\lambda_j$ are parameters that account for the hydrophobicity of the $i$th and $j$th interacting particles respectively, being $\lambda_{ij} = (\lambda_i + \lambda_j)/2$. The excluded volume of the different residues/nucleotides is given by $\sigma_i$ and $\sigma_j$, where $\sigma_{ij}$ = $(\sigma_i + \sigma_j)/2$, and $r$ is the distance between the $i$th and $j$th particles. $\epsilon_{ij}$ (0.2 kcal/mol) is a fitting parameter to reproduce the experimental single-IDR radius of gyration ($R_g$) [18]. When at least one of the $i$th or $j$th amino acids is part of a structured globular domain, $\lambda_{ij}$ is scaled by a factor of 0.7 to account for 'buried' amino acids in globular domains [20]. The specific values for each amino acid and nucleotide parameters $\sigma$, $q$, and $\lambda$ can be found in References: Dignon *et al.* for FUS [18] and Regy *et al.* for RNA [19].

Finally, we consider an additional term to describe cation-$\pi$ (c-$\pi$) interactions (only for the following set of pairs of amino acids (c-$\pi$:{Arg-Phe, Arg-Trp, Arg-Tyr, Lys-Phe, Lys-Trp and Lys-Tyr}):

$$E_{\text{cation-}\pi} = \sum_{i \in \{\text{c-}\pi\}} \sum_{\substack{j \in \{\text{c-}\pi\} \\ j<i}} 4\epsilon_{ij} \left[ \left(\frac{\sigma_{ij}}{r}\right)^{12} - \left(\frac{\sigma_{ij}}{r}\right)^{6} \right], \tag{S5}$$

where $\sigma_{ij}$ is the same as in the hydrophobic interactions, and $\epsilon_{ij}$ is 3.0 kcal mol$^{-1}$ for all six cation-$\pi$ pairs as proposed in Ref. [17] (Approach 1). Consistently, the interaction of these amino acids is scaled down by a 30% when they are found in structured globular domains.

We carry out Direct Coexistence (DC) simulations [12] using 48 protein replicas to compute phase diagrams in the NVT ensemble (*i.e.*, constant number of particles (N), volume (V) and temperature (T)). For all our simulations with this force field, we use the LAMMPS MD package [23]. Since FUS, reordered FUS, and mutated FUS include globular domains, we perform simulations using the LAMMPS rigid body Nosé–Hoover thermostat [14–16] in combination with a Langevin thermostat [24] for the non-rigid bodies (both with a relaxation time of 5 ps). We apply periodic boundary conditions in the three directions of space. A time step of 10 fs has been employed for the Verlet integration of the equations of motion. LARKS disorder-to-order transitions are implemented through our dynamical ageing model in bulk phases at condensate coexistence densities (derived from DC simulations) running NVT simulations. We run two independent trajectories for each studied system (*i.e.*, original, mutated, and reordered FUS variants) in order to reduce the uncertainty in our simulations. Hence, the corresponding results of this model shown in the main text are always obtained by averaging the number of disorder-to-order transitions over time using two different independent simulations. Viscoelastic calculations (*i.e.*, $G(t)$) have been evaluated throughout the cubic NVT simulations. The volume of each box is set to the corresponding bulk equilibrium density of condensate at the temperature of interest. The typical length of each simulation to compute viscosity is about 5 $\mu$s.

#### D. Protein sequences and PDB of the structured domains

**FUS**

MASNDYTQQATQSYGAYPTQPGQGYSQQSSQPYGQQSYSGYSQSTDTSGYGQSSYSSYGQSQNTGYGTQSTPQGYGSTGGYGSS
QSSQSSYGQQSSYPGYGQQPAPSSTSGSYGSSSQSSSYGQPQSGSYSQQPSYGGQQQSYGQQQSYNPPQGYGQQNQYNSSSGGGG
GGGGGGNYGQDQSSMSSGGGSGGGYGNQDQSGGGGSGGYGQQDRGGRGRGGSGGGGGGGGGGYNRSSGGYEPRGRGGGRG
GRGGMGGSDRGGFNKFGGPRDQGSRHDSEQDNSDNNTIFVQGLGENVTIESVADYFKQIGIIKTNKKTGQPMINLYTDRETGKL
KGEATVSFDDPPSAKAAIDWFDGKEFSGNPIKVSFATRRADFNRGGGNGRGGRGRGGPMGRGGYGGGGSGGGGRGGFPSGGG
GGGGQQRAGDWKCPNPTCENMNFSWRNECNQCKAPKPDGPGGGPGGSHMGGNYGDDRRGGRGGYDRGGYRGRGGDRGGF
RGGRGGGDRGGFGPGKMDSRGEHRQDRRERPY

The following Protein Data Bank (PDB) codes were used to build the structured globular domains of FUS (residues from 285–371 (PDB code: 2LCW) and from 422–453 (PDB code: 6G99)).

**Mutated-FUS**

MASNDYTQQATQSYGAYPTQPGQGYSQQRRQPYRQQSYSGYSQSTDTRRYGQSSYSSYGQSQNTRYRTQSTPQGYGSTGGYGS
SQRRQSSYRQQSSYPGYGQQPAPSSTSGSYGSSSQSSSYGQPQSGSYSQQPSYGGQQQSYGQQQSYNPPQGYGQQNQYNSSSGGG
GGGGGGGNYGQDQSSMSSGGGSGGGYGNQDQSGGGGSGGYGQQDRGGRGRGGSGGGGGGGGGGYNRSSGGYEPRGRGGGR
GGRGGMGGSDRGGFNKFGGPRDQGSRHDSEQDNSDNNTIFVQGLGENVTIESVADYFKQIGIIKTNKKTGQPMINLYTDRETGK
LKGEATVSFDDPPSAKAAIDWFDGKEFSGNPIKVSFATRRADFNRGGGNGRGGRGRGGPMGRGGYGGGGSGGGGRGGFPSGG
GGGGGQQRAGDWKCPNPTCENMNFSWRNECNQCKAPKPDGPGGGPGGSHMGGNYGDDRRGGRGGYDRGGYRGRGGDRGG
FRGGRGGGDRGGFGPGKMDSRGEHRQDRRERPY

**Reordered-FUS**

GPRDQGSRHDSEQDNSDNNTIFVQGLGENVTIESVADYFKQIGIIKTNKKTGQPMINLYTDRETGKLKGEATVSFDDPPSAKAAID
WFDGKEFSGNPIKVSFATRSSSGGGGGGGGGGNYGQDQSSMSSGGGSGGGYGNQDQSGGGGSGGYGQQDRGGRGRGGSGGG
GGGGGGGYNRSSGGYEPRGRGGGRGGRGGMGGSDRGGFNKFGMASNDYTQQATQSYGAYPTQPGQGYSQQSSQPYGQQSYS
GYSQSTDTSGYGQSSYSSYGQSQNTGYGTQSTPQGYGSTGGYGSSQSSQSSYGQQSSYPGYGQQPAPSSTSGSYGSSSQSSSYGQP
QSGSYSQQPSYGGQQQSYGQQQSYNPPQGYGQQNQYNRADFNRGGGNGRGGRGRGGPMGRGGYGGGGSGGGGRGGFPSGG
GGGGGQQRAGDWKCPNPTCENMNFSWRNECNQCKAPKPDGPGGGPGGSHMGGNYGDDRRGGRGGYDRGGYRGRGGDRGG
FRGGRGGGDRGGFGPGKMDSRGEHRQDRRERPY

While for the mutated variant, the structured globular domains correspond to the same residues specified for the original sequence, for the reordered variants these domains will correspond to the following regions: residues from 19-105 (PDB code: 2LCW) and from 422–453 (PDB code: 6G99)).

For hnRNPA1 the structured domains correspond to the residues from 8-91 and 103-181 (both taken from PDB code: 1L3K).

**hnRNPA1**

MSKSESPKEPEQLRKLFIGGLSFETTDESLRSHFEQWGTLTDCVVMRDPNTKRSRGFGFVTYATVEEVDAAMNARPHKVDGRV
VEPKRAVSREDSQRPGAHLTVKKIFVGGIKEDTEEHHLRDYFEQYGKIEVIEIMTDRGSGKKRGFAFVTFDDHDSVDKIVIQKYH
TVNGHNCEVRKALSKQEMASASSSQRGRSGSGNFGGGRGGGFGGNDNFGRGGNFSGRGGFGGSRGGGGYGGSGDGYNGFGND
GGYGGGGPGYSGGSRGYGSGGQGYGNQGSGYGGSGSYDSYNNGGGGGFGGGSGSNFGGGGSYNDFGNYNNQSSNFGPMKGG
NFGGRSSGPYGGGGQYFAKPRNQGGYGGSSSSSSYGSGRRF

**Reordered hnRNPA1**

For reordered hnRNPA1 the structured domains correspond to the residues from 8-91 and 235-313 (both taken from PDB code: 1L3K).

MSKSESPKEPEQLRKLFIGGLSFETTDESLRSHFEQWGTLTDCVVMRDPNTKRSRGFGFVTYATVEEVDAAMNARPHKVDGRV
VEPKRAVSREDSQRPGAHLGDGYNGFGNDGGYGGGGPGYSGGSRGYGSGGQGYGNQGSGYGGSGSYDSYNNGGGGGFGGGS
GSNFGGGGSYNDFGNYNNQSSNFGPMKGGNFGGRSSGPYGGGGQYFAKPRNQGGYGGSSSSSSYGSGRRFTVKKIFVGGIKEDT
EEHHLRDYFEQYGKIEVIEIMTDRGSGKKRGFAFVTFDDHDSVDKIVIQKYHTVNGHNCEVRKALSKQEMASASSSQRGRSGSGN
FGGGRGGGFGGNDNFGRGGNFSGRGGFGGSRGGGGYGGS

### SII. OBTAINING PHASE DIAGRAMS VIA DIRECT COEXISTENCE SIMULATIONS

To obtain the phase diagrams presented in this work (Figs. 2(a), 3(b), and 4(a) of the main text), we compute the coexistence density of each phase at different temperatures employing the Direct Coexistence method [12, 25, 26]. We put in contact in the same simulation box the two coexisting protein/RNA phases: the condensed and the diluted phase. Once the system reaches equilibrium, we compute the density profile along the long axis of the box, and thus, we extract the density of the two coexisting phases (*i.e.*, Fig. 2(a) and Fig. 3(b) of the main text). The critical point of the phase diagrams is estimated by using the universal scaling law of coexistence densities near a critical point [27], and the law of rectilinear diameters [28]:

$$\big(\rho_l(T) - \rho_v(T)\big)^{3.06} = d\left(1 - \frac{T}{T_c}\right), \tag{S6}$$

and

$$(\rho_l(T) + \rho_v(T))/2 = \rho_c + s_2(T_c - T), \tag{S7}$$

where $\rho_v$ and $\rho_l$ are the coexisting densities of the diluted and condensed phases, respectively, $T_c$ and $\rho_c$ are the critical temperature and density, respectively, and $d$ and $s_2$ are the fitting parameters. The critical temperatures of the different proteins studied prior-ageing (which roughly coincides with that post-ageing) are provided in Table S2.

| LCD | FUS | Mutated FUS | Reordered FUS |
|---|---|---|---|
| $(329 \pm 5)\,K$ | $(396 \pm 5)\,K$ | $(389 \pm 5)\,K$ | $(405 \pm 5)\,K$ |

**TABLE S2:** Critical temperature of the different FUS sequences and the different LCD variants prior-ageing. Please note that the critical temperature for all LCD sequences prior-ageing is the same by construction (329K) since the interactions between non-structured (LARKS or non-LARKS) residues is the same.

In Fig. 4(a) of the main text, we show the critical temperature of the different FUS sequences as a function of the polyU/FUS mass ratio. To determine these critical temperatures, in Fig. S1, we present the phase diagrams (in the $T - \rho$ plane) for the reordered and mutated FUS variants at increasing concentrations of polyU RNA (of 125 nucleotide strands). The phase diagram for the original FUS sequence as a function of polyU RNA (of the same strand length) can be found in Ref. [29].

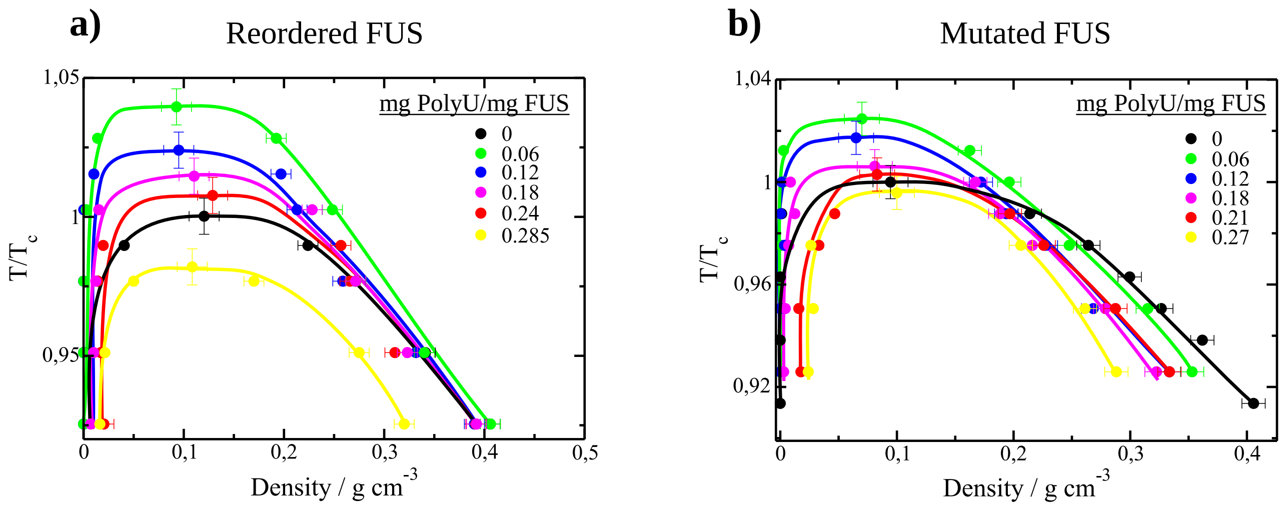


**FIG. S1:** Phase diagram in the $T-\rho$ plane for the two studied FUS variants at different concentrations of polyU RNA (expressed in mass ratio as indicated in the legend). Temperature has been renormalized by the critical temperature of each FUS variant in absence of RNA ($T_c$; provided in Table S2). a) Reordered FUS variant. b) Mutated FUS variant.

### SIII. AGEING DYNAMIC ALGORITHM

In this work we have characterised the condensate ageing of different archetypal LCD sequences as well as of distinct FUS sequences due to disorder-to-order structural transitions. For that purpose, we have employed a time and local-dependent dynamic algorithm based on our previous work from Ref. [30]. Through this algorithm, the Low-complexity Aromatic-Rich Kinked Segments (LARKS) of FUS, such as $_{37}$SYSGYS$_{42}$, $_{54}$SYSSYGQS$_{61}$, and $_{77}$STGGYG$_{82}$, as well as those proposed for the different studied LCDs can undergo 'effective' structural transitions from disordered states to inter-protein $\beta$-sheets (in terms of binding energy and structure) depending on the local environment. To recapitulate in our simulations such transitions, every 100 simulation timesteps, if the conditions around each of the disordered LARKS are favourable to form a cross-$\beta$-sheet motif (*i.e.*, a high-density fluctuation), the dynamic algorithm changes the force field parameters to those given in Table S3 (obtained from atomistic simulations; Fig. 1 of the main text) corresponding to inter-protein structured $\beta$-sheets. The dynamical evaluation of these conditions takes place on the central bead of each LARKS (the chosen amino acid for which we perform the calculations is detailed in Table S3). We employ a cut-off distance criterion ($r_{cut}$ in Table S3) to evaluate the required conditions and trigger the structural $\beta$-sheet transitions. This occurs when four LARKS met within a given cut-off distance (Table S3) [30]. Then, the average effective interaction binding between these LARKS is replaced by that obtained from atomistic PMF simulations (*i.e.*, the $\lambda$ of the corresponding amino acids of a given LARKS is updated to a global $\lambda_{ordered}$ provided in Tables S3 and S4 for the whole LARKS sequence once the transition has taken place). For the case of the different archetypal LCD sequences (Table S4), the effective interaction between the structured LARKS has been obtained by averaging the energy of the structured LARKS of the sequences shown in Figure 1(b) of the main text. Importantly, cross-$\beta$-sheet transitions can be reversible. If one of the structured LARKS separates from the cross-$\beta$-sheet motif beyond a given distance ($r_{cut,reverse}$ in Tables S3 and S4), the interaction parameters are reversed to those of the disorder residues (HPS-Cation-$\pi$ force field). After a structural transition takes place, we assign an average mass to the new (averaged) structured LARKS residues which keeps constant the total mass of the proteins. Furthermore, to recapitulate the local increase of rigidity associated with a $\beta$-sheet structural transition, we introduce an angular term to the total energy (Equation S1) that follows the following equation:

$$E_{\text{Angles}} = \sum_{\text{Angles}} k_{ang}(\theta - \theta_0)^2, \tag{S8}$$

where we set $\theta_0 = 180°$ and $k_{ang} = 5$ kcal mol$^{-1}$ rad$^{-2}$ for the structured LARKS of an inter-protein $\beta$-sheet. To carry out these simulations, we use the USER-REACTION [31] package of LAMMPS which allows us to change the topology of the underlying system components in a time-dependent and local-dependent manner.

| | FUS (1) | FUS (2) | FUS(3) |
|---|---|---|---|
| Central amino acid | $S_{39}$ | $S_{57}$ | $G_{80}$ |
| $m_{ordered}$ / (g mol$^{-1}$) | 107.44 | 107.48 | 86.92 |
| $\lambda_{ordered}$ / (kcal mol$^{-1}$) | 2.8 | 3.5 | 2.2 |
| $\sigma_{ordered}$ / Å | 5.52 | 5.52 | 5.13 |
| $r_{cut}$ / Å | 8.2 | 8.2 | 8.3 |
| $r_{cut,reverse}$ / Å | - | - | 20 |

**TABLE S3:** Set of parameters employed for residues belonging to structured inter-peptide $\beta$-sheet motifs in FUS. FUS (1), FUS (2) and FUS (3) correspond to the $_{37}$SYSGYS$_{42}$, $_{54}$SYSSYGQS$_{61}$ and $_{77}$STGGYG$_{82}$ sequences respectively. The central amino acid indicates the residue used by the ageing algorithm to evaluate LARKS high-density fluctuations. $m_{ordered}$, $\lambda_{ordered}$, and $\sigma_{ordered}$ refer to the mass, interaction energy, and molecular diameter respectively of the residues composing the structured LARKS of each sequence. Please note that in our model, after the structural transition takes place, the different residues of a given sequence are replaced by an average mutated residue that conserves the volume and mass of the unstructured LARKS. Finally, $r_{cut}$ and $r_{cut,reverse}$ account for the cut-off distances employed for triggering and reversing, respectively, the structural transitions depending on the LARKS environment. Please note that since in FUS(1) and FUS(2) the formation of cross-$\beta$-sheets is almost irreversible, we do not include back transitions to speed up our simulations.

| | LCD |
|---|---|
| $m_{ordered}$ / (g mol$^{-1}$) | 118.9 |
| $\epsilon_{ordered}$ / (kcal mol$^{-1}$) | 1.53 |
| $\sigma_{ordered}$ / Å | 5.25 |
| $r_{cut}$ / Å | 6.9 |
| $r_{cut,reverse}$ / Å | 10.5 |

**TABLE S4:** Set of parameters employed for residues belonging to structured inter-peptide $\beta$-sheet motifs within LCD sequences. The central amino acid employed by the ageing algorithm to evaluate LARKS high-density fluctuations changes with the sequence as follows: 1-Tail (7), 1-Center(62), 2-SameTail (7,27), 2-OppositeTail (7,118), 2-Center (52,72), 2-Equispaced (40,83), 3-Equispaced (29,61,93). $m_{ordered}$, $\lambda_{ordered}$, and $\sigma_{ordered}$ refer to the mass, interaction energy, and molecular diameter respectively of the residues composing the structured LARKS of each sequence. Finally, $r_{cut}$ and $r_{cut,reverse}$ account for the cut-off distances employed for triggering and reversing, respectively, the structural transitions depending on the LARKS local environment.

The criterion that at least four peptides should be in close contact to trigger a structural disorder-to-order transition has been chosen based on the following reasons. Four interacting peptides is the minimal system in which all the different types of stacking and hydrogen bonding interactions that stabilise the $\beta$-sheet fibrillar ladder are fulfilled (*i.e.*, the two interacting steps of the ladder are formed each by a pair of $\beta$-sheet peptides). Thus, considering fewer interacting peptides (e.g. only two or three) would severely underestimate the strength of interactions among ordered LARKS in our atomistic simulations and, subsequently, erroneously preclude the formation of kinetically arrested states. Atomistic calculations from previous studies (Refs. [11, 30, 32]) show that the strength of interactions among four structured peptides is already high enough for the LARKS to remain stably bound upon thermal fluctuations. Hence, if we made the criterion even more stringent (*i.e.*, by requiring clustering of five or more peptides, the strength of interactions among the system would remain consistent with kinetic arrest at the atomistic level (as shown in Fig. 1(a) of the main text). However, such more stringent criterion would now render coarse-grained simulations prohibitively expensive. That is, much longer simulation timescales would be needed to capture the rarer higher-density fluctuations that could result in the spontaneous formation of clusters of five or more peptides. This would be opposed to the more frequent fluctuations that yield clusters of four peptides, which are already strong enough to maintain $\beta$-sheet clusters stably bound, as shown in Ref. [30]. In that sense, despite fluctuations of 4 or more LARKS can also trigger a $\beta$-sheet transition in our model, a threshold of at least four peptides (rather than two or three) is set to guarantee long-lived binding [11, 30, 32]. Moreover, since experimental $\beta$-sheet transition barriers suggest typical timescales of the order of hundreds of nanoseconds [33–35], and protein self-diffusion timescales are of the order of hundreds of milliseconds [36], we approximate in our ageing dynamic algorithm (following Refs. [30, 32]) a negligible transition barrier once protein density fluctuations bring together four LARKS in close contact (i.e., within the given cut-off distances provided in Tables S3 and S4).

### SIV. ANALYSIS OF THE NETWORK CONNECTIVITY BY A PRIMITIVE PATH ANALYSIS

We have analyzed the $\beta$-sheet network connectivity of the aged condensates by means of the primitive path analysis (PPA) method [37]. These calculations minimize the contour length of all chains in the system, while keeping the end monomers fixed, and thus respecting the network topology by not allowing chains to cross each other. We have employed in this case a recently modification of this algorithm proposed by Tejedor *et al.* [32]. For this purpose, we first perform NpT simulations to allow the number of cross-$\beta$-sheet transitions in each of the different studied proteins to reach a steady state. Within the employed modification, we freeze the positions of the inter-protein $\beta$-sheet motifs. Then, we perform an energy minimization in which the bond length is fix to zero and where the excluded volume interactions are removed. Thus, the contour length of the strands between the transitioned structured LARKS is minimized while preserving the underlying network connectivity. To run the energy minimizations required for the PPA method, we have used LAMMPS [13] using $1 \cdot 10^5$ iterations with a tolerance of $1 \cdot 10^{-6}$ for both the energy and the total force.

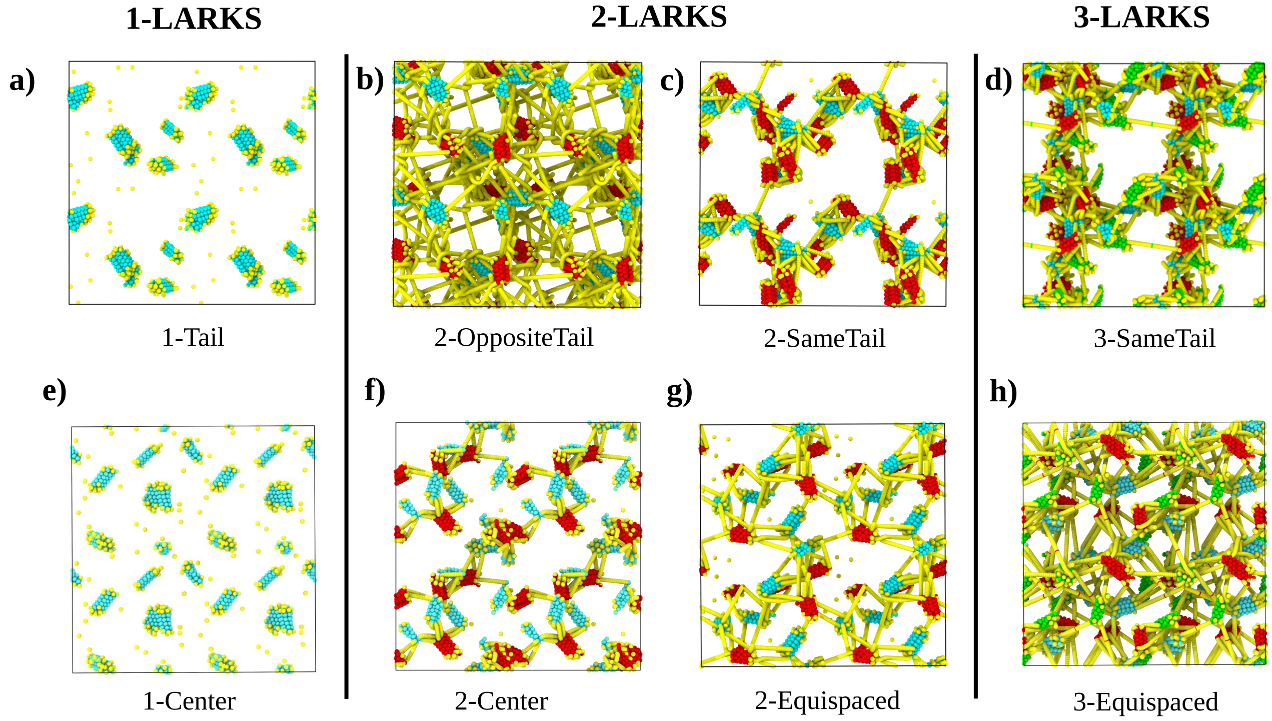


**FIG. S2:** Network connectivity of low-complexity domains with different LARKS abundance and patterning evaluated using the primitive path analysis after 100 ns of maturation at $T/T_c = 0.91$ in a cubic box at the corresponding condensate bulk density for such temperature: a) 1-Tail. b) 2-OppositeTail. c) 2-SameTail. d) 3-SameTail. e) 1-Center. f) 2-Center. g) 2-Equispaced. h) 3-Equispaced.

Figure S2 shows the network connectivity of the aged LCD condensates proposed in this work. As we point out in the main text, only the LCDs with 1 LARKS are able to relax (Fig. 2(c) main text) as we can observe by the network connectivity analysis. For the rest of LCD sequences, condensates are not able to relax since they show a strong network of inter-protein $\beta$-sheet contacts which is fully percolated. In Figure S3, the network connectivity of the three FUS sequences upon maturation, both in absence and presence of RNA, are shown.

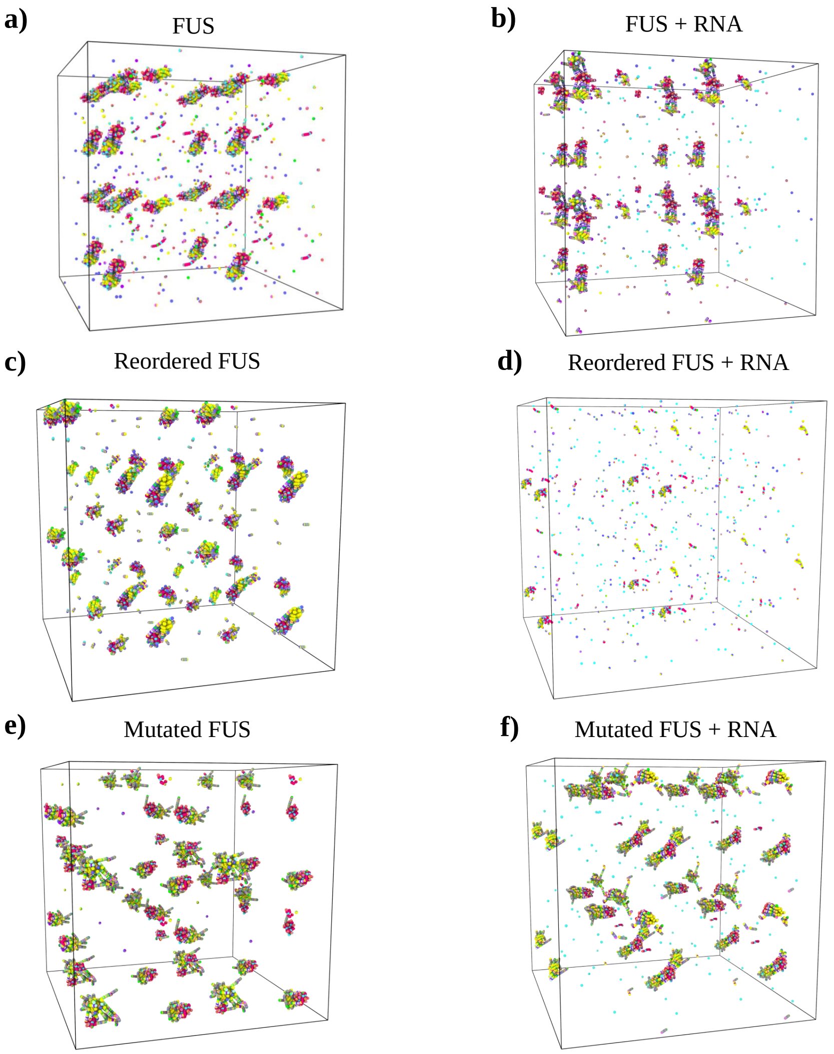


**FIG. S3:** Network connectivity of the different FUS sequences calculated with the PPA method after 300 ns of maturation at $T/T_{c,pure}$ ∼0.97 within a cubic box at the bulk condensate density at such temperature: a) FUS b) FUS in presence of 0.24 mg of polyU per mg of protein. c) Reordered FUS variant. d) Reordered FUS variant in presence of 0.285 mg of polyU per mg of protein. e) Mutated FUS variant. f) Mutated FUS variant in presence of 0.27 mg of polyU per mg of protein.

### SV. TIME-EVOLUTION OF INTER-PROTEIN $\beta$-SHEET TRANSITIONS THROUGH A SECOND-ORDER REACTION ANALYSIS.

To quantify the progressive formation of disorder-to-order transitions within condensates of the different FUS sequences (Figs. 3 and 4 of the main text) and the proposed low-complexity domains (Figs. 1 and 2 of the main text), we apply a second-order reaction analysis. We start by defining $[D]$ as the concentration of reactive sites in their disordered state, and $[S]$ as the concentration of structured $\beta$-sheet domains (please note that when a sequence contains more than one LARKS, we average the number of reactions per number of LARKS). Then, we assume that the reactions between $D$ and $S$ follow the next mechanism:

$$\begin{aligned} 2D &\xrightarrow{k_{dir}} 2S \\ S + D &\xrightarrow{k_{dir}} 2S \\ D &\xleftarrow{k_{inv}} S, \end{aligned} \tag{S9}$$

where $k_{dir}$ is the kinetic constant for the direct reaction, and $k_{inv}$ is the kinetic constant for the inverse reaction.

If we assume that there is a complete mixing of all the species of the system and we define the equilibrium constant as $K_{eq} = \frac{k_{dir}}{k_{inv}}$, we can obtain the following system of ordinary differential equations for the time evolution of the concentrations of $D$ and $S$:

$$\begin{aligned} \frac{d[D]}{dt} &= -2k_{dir}[D]^2 - k_{dir}[D][S] + \frac{k_{dir}}{K_{eq}}[S] \\ \frac{d[S]}{dt} &= 2k_{dir}[D]^2 + k_{dir}[D][S] - \frac{k_{dir}}{K_{eq}}[S]. \end{aligned} \tag{S10}$$

This system can be solved employing an adaptive time step algorithm for different values of the parameters $k_{dir}$ and $K_{eq}$, and then used for fitting the time-evolution of the disorder-to-order transitions. Hence, the obtained fitting parameters correspond to the kinetic constants. In the main text, we show the results of $k_{dir}$ for every LCD sequence proposed (Fig. 2(b)), and the values of $k_{dir}$ as function of the RNA concentration for FUS, mutated FUS and reordered FUS (Figs. 3(c) and 4(c)). Here, we provide the numerical values of both constants $k_{dir}$ and $K_{eq}$ collected in Table S5 for FUS, mutated FUS and reordered FUS/polyU mixtures. Furthermore, in Tables S6, S7, and S8, we present the obtained results for the distinct LCD sequences. Finally we have collected the results for hnRNPA1 in Table S9 Please note that the error in the estimates of $K_{eq}$ may be specially significant in some cases due to the fact that transitions are mostly irreversible under some of the studied conditions, and (2) the steady state is not well-captured in some condensates possibly yielding misleading values of the equilibrium constant $K_{eq}$.

| FUS | | | Reordered FUS | | | Mutated FUS | | |
|---|---|---|---|---|---|---|---|---|
| RNA/protein mass ratio | $K_{eq}$ | $k_{dir}$ $(ms^{-1}\%^{-1})$ | RNA/protein mass ratio | $K_{eq}$ | $k_{dir}$ $(ms^{-1}\%^{-1})$ | RNA/protein mass ratio | $K_{eq}$ | $k_{dir}$ $(ms^{-1}\%^{-1})$ |
| 0.00 | 7.3e+04 ± 7.7 | 12.409 ± 0.019 | 0.00 | 1.07e+06 ± 1.31 | 9.37 ± 0.02 | 0.00 | 3.25 ± 0.06 | 41.9 ± 0.1 |
| 0.12 | 1.5e+05 ± 1.9 | 12.371 ± 0.020 | 0.12 | 1.60 ± 0.08 | 11.80 ± 0.04 | 0.12 | 2.85e+06 ± 7.85 | 34.50 ± 0.07 |
| 0.24 | 2.1 ± 0.054 | 11.532 ± 0.023 | 0.24 | 55 ± 7.6 | 1.350 ± 0.004 | 0.24 | 1.39e+07 ± 2.77 | 34.40 ± 0.09 |
| | | | 0.285 | 6.7e+03 ± 1.05 | 0.65 ± 0.015 | 0.27 | 3.47e+07 ± 1.78 | 25.1 ± 0.3 |

**TABLE S5:** Kinetic constants obtained from fits to a second-order reaction model for FUS, Reordered FUS and Mutated FUS/polyU bulk condensates using the HPS-Cation−$\pi$ force field, and at $T/T_{c,pure}$ ∼0.97 and different polyU RNA/protein mass ratios.

| 1-Tail | | 1-Center | |
|---|---|---|---|
| $K_{eq}$ | $k_{dir}$ $(ms^{-1}\%^{-1})$ | $K_{eq}$ | $k_{dir}$ $(ms^{-1}\%^{-1})$ |
| 3.56e+08 ± 1.45 | 109.0 ± 0.1 | 6.48e+07 ± 6.35 | 65.7 ± 0.3 |

**TABLE S6:** Kinetic constants obtained from fits to a second-order reaction model for 1-LARKS LCD bulk condensates at $T/T_c$=0.953 and their corresponding density at such temperature (Fig. 2(a) of the main text).

| 2-Center | | 2-Equispaced | | 2-SameTail | | 2-OppositeTail | |
|---|---|---|---|---|---|---|---|
| $K_{eq}$ | $k_{dir}$ $(ms^{-1}\%^{-1})$ | $K_{eq}$ | $k_{dir}$ $(ms^{-1}\%^{-1})$ | $K_{eq}$ | $k_{dir}$ $(ms^{-1}\%^{-1})$ | $K_{eq}$ | $k_{dir}$ $(ms^{-1}\%^{-1})$ |
| -3.74e+08 ± 5.55 | 63.0 ± 0.2 | 8.09e+07 ± 5.0 | 66.4 ± 0.1 | -7.48e+07 ± 1.0 | 113.0 ± 0.4 | -1.87e+08 ± 3.77 | 119.0 ± 0.2 |

**TABLE S7:** Kinetic constants obtained from fits to a second-order reaction model for 2-LARKS LCD bulk condensates at $T/T_c$=0.953 and their corresponding density at such temperature (Fig. 2(a) of the main text).

| 3-Equispaced | | 3-SameTail | |
|---|---|---|---|
| $K_{eq}$ | $k_{dir}$ $(ms^{-1}\%^{-1})$ | $K_{eq}$ | $k_{dir}$ $(ms^{-1}\%^{-1})$ |
| 6.23e+08 ± 8.44 | 69.8 ± 0.2 | 6.63e+06 ± 1.42 | 93.1 ± 0.7 |

**TABLE S8:** Kinetic constants obtained from fits to a second-order reaction model for 3-LARKS LCD bulk condensates at $T/T_c$=0.953 and their corresponding density at such temperature (Fig. 2(a) of the main text).

| hnRNPA1 | | Reordered hnRNPA1 | |
|---|---|---|---|
| $K_{eq}$ | $k_{dir}$ $(ms^{-1}\%^{-1})$ | $K_{eq}$ | $k_{dir}$ $(ms^{-1}\%^{-1})$ |
| 3.06e+06 ± 3.97 | 18.02 ± 0.01 | 2.2e+03 ± 9.6 | 8.25 ± 0.03 |

**TABLE S9:** Kinetic constants obtained from fits to a second-order reaction model for hnRNPA1 and reordered hnRNPA1 bulk condensates using the HPS-Cation$-\pi$ force field, and at $T/T_{c,pure}$ ~0.97; referring $T_c$ to the critical temperature of each sequence (see phase diagram in Fig. S9(b)).

### SVI. CALCULATION OF MOLECULAR CONTACT PROBABILITIES

Landscapes of intermolecular protein contact probabilities are calculated throughout NVT simulations at constant temperature and equilibrium bulk density. These results are obtained from the same simulations employed to calculate the transport properties within protein condensates (see Section SVII) of the studied systems (*i.e.*, FUS, reordered FUS, mutated FUS, and the different LCD sequences). The contact maps are calculated from n=2 independent trajectories which contain ~150 independent configurations. We have used the 'smart' cut-off analysis introduced in our recent work [29]. In that analysis, the cut-off distance is set to $1.2\sigma_{ij}$, where $\sigma_{ij}$ accounts for the mean excluded volume of each specific pair $i$th and $j$th amino acids. Since the minimum of the pairwise potential (Eqs. (S4) and (S5)) is located at $2^{1/6}\sigma_{ij} \approx 1.122\sigma_{ij}$, we set our cut-off distance slightly beyond that point, at $1.2\sigma_{ij}$, to ensure significant attractive binding between pairs of amino acids.

The intermolecular contact map probability estimates the average number of attractive interactions—sustaining protein aggregation—that each residue along the protein sequence establishes within bulk phase-separated droplets per protein replica. Thus, we can compare the maps of the same system from both pre-aged and aged-condensates (after a certain time, which is 100 ns in the case of the LCDs and 300 ns in the case of FUS sequences) to explore the effect of accumulation of inter-protein $\beta$-sheets motifs in matured condensates. From these probabilities, we can infer a binding free energy variation (if assuming that aged condensates at extremely long timescales can be still equilibrated):

$$\frac{\Delta G}{k_B T} \approx -ln\left(\frac{P_{post}}{P_{pre}}\right), \tag{S11}$$

where $P_{pre}$ and $P_{post}$ correspond to residue contact probabilities for pre-aged and post-aged amino acid pairs respectively, $T$ is the temperature, and $k_B$ is the Boltzmann constant. It is worth noting that the calculation using Eq. (S11) can be flawed if any residue-residue contact never happens, either in the numerator or the denominator inside the logarithm. To sort out such numerical drawback, we rescale both probabilities by summing one contact to every element of both maps so the divergent points are reduced to a reasonable value.

In this work, we have calculated the binding probability difference contact maps of the distinct LCD sequences upon maturation (Fig. 2(e) of the main text shows the map for the 3-Equispaced LCD patterning). To evaluate the free energy difference between pre and post aged condensates, we calculate first the maps of intermolecular contact residues per amino acid of the distinct LCD condensates, which are shown in Fig. S4 for the aged condensates. Once we have calculated these maps, we can compute the difference in contact probability upon maturation, as shown in Fig. S5). For the different FUS sequences (in absence of polyU RNA), the same information is provided in Figs. S6 and S7.

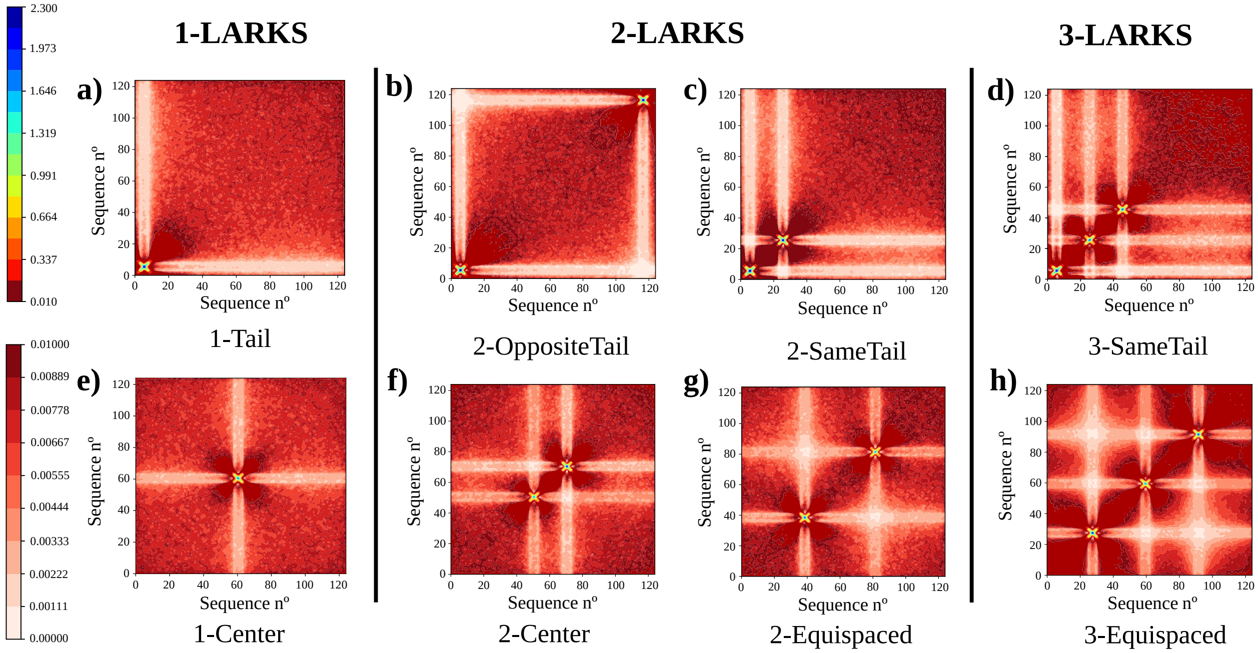


**FIG. S4:** Average number of intermolecular contact residues per amino acid for the different LCD sequences calculated through the 'smart cut-off' distance analysis within phase-separated droplets after 100 ns of maturation, at $T/T_c = 0.91$, and using a cubic box with the bulk condensate density: a) 1-Tail. b) 2-OppositeTail. c) 2-SameTail. d) 3-SameTail e) 1-Center. f) 2-Center. g) 2-Equispaced. h) 3-Equispaced. Please note that two different colour scales have been included to improve the visualization of the intermolecular contact maps.

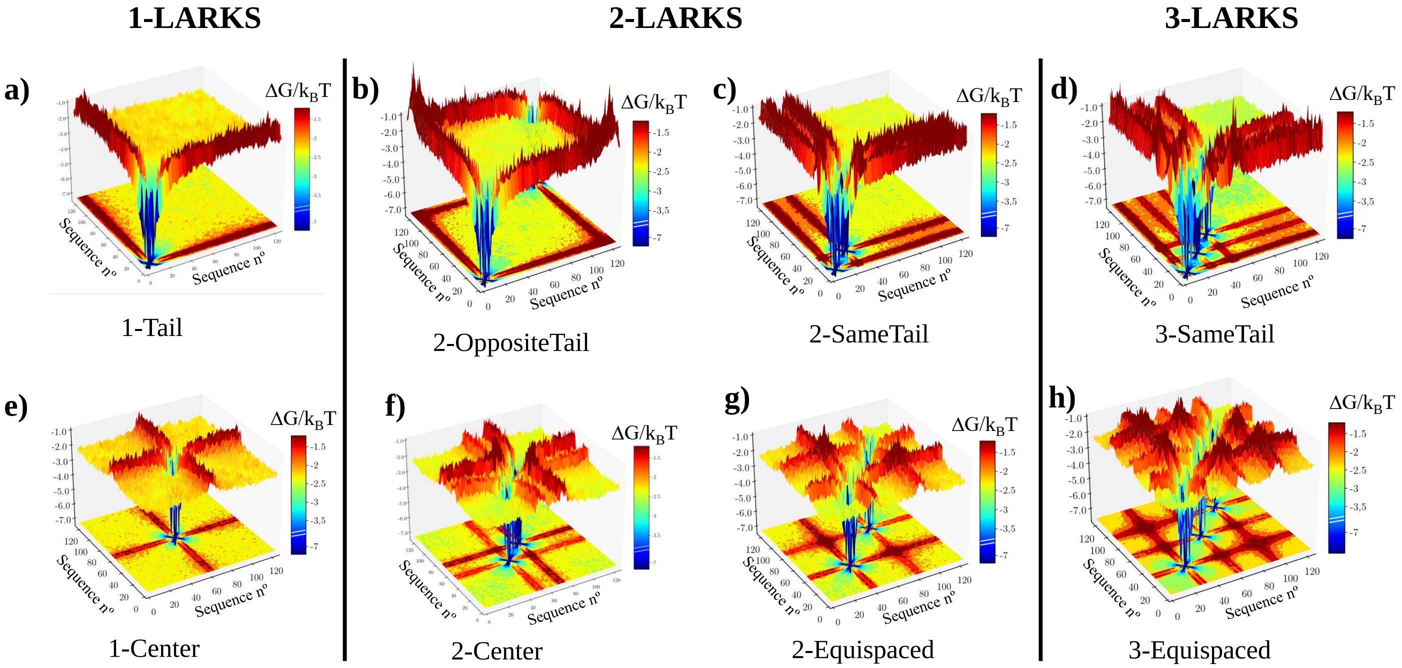


**FIG. S5:** Landscape of the protein contact free energy variation upon condensate maturation (100 ns of ageing, $T/T_c = 0.91$, and bulk condensate density) for the different LCD sequences. $\Delta G/k_BT$ is obtained from the residue contact probability ratio between aged condensates and liquid-like condensates before ageing. Colour map projections of the free energy landscape in 2-dimensions are also included.

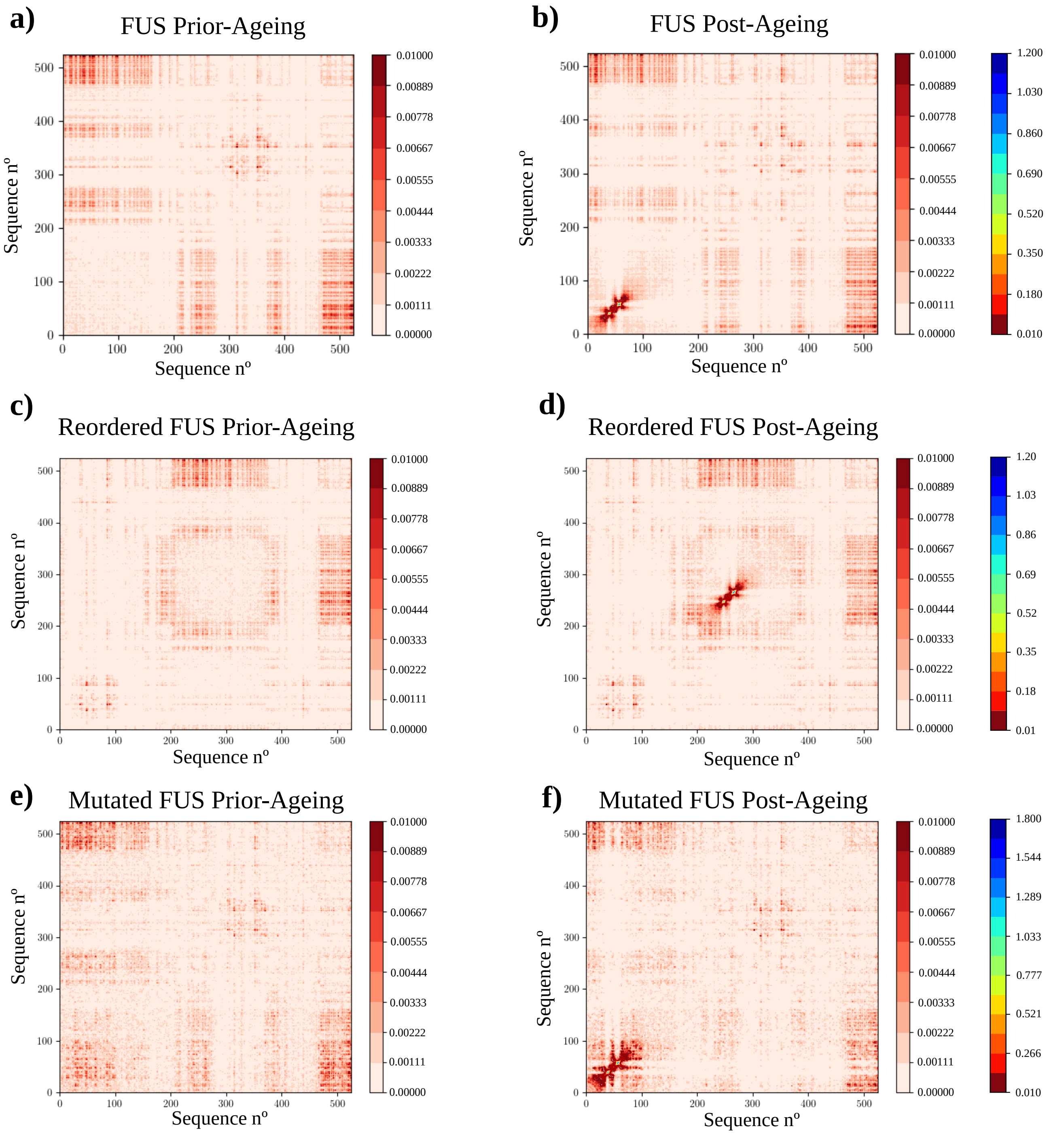


**FIG. S6:** Average number of intermolecular contact residues per amino acid for the different FUS sequences calculated through the 'smart cut-off' distance analysis within phase-separated droplets prior-ageing and after 300 ns of maturation at $T/T_{c,pure} \sim 0.97$, and at the bulk condensate density. Please note that two different colour scales have been included to improve the visualization of the intermolecular contact maps.

### SVII. CALCULATION OF TRANSPORT PROPERTIES WITHIN PHASE-SEPARATED CONDENSATES

The calculation of viscosity for phase-separated condensates has been done by running NVT simulations of a cubic box with a fixed volume corresponding to the condensate bulk density. We have measured the viscosity of the different

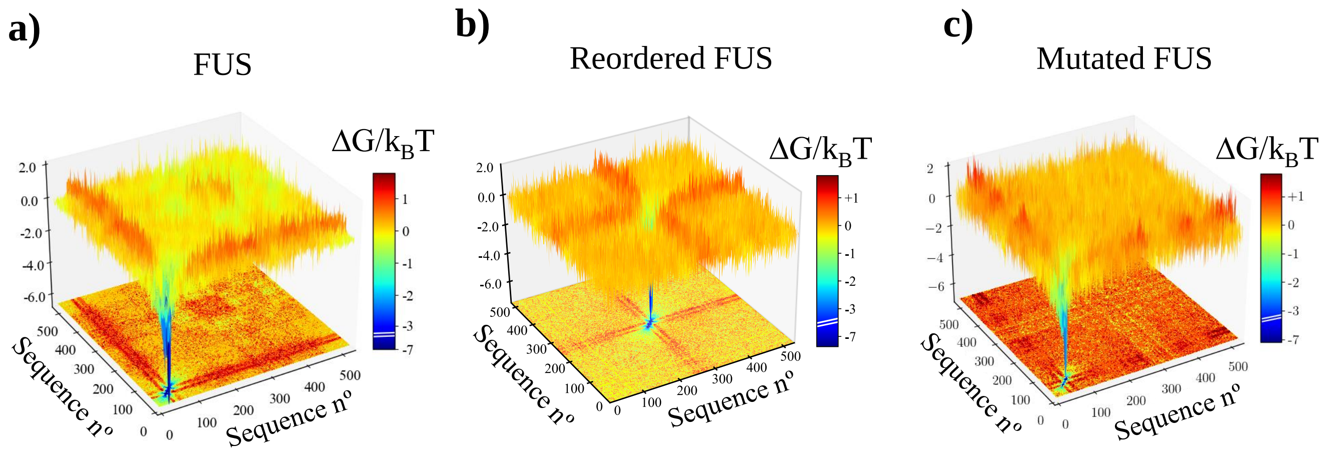


**FIG. S7:** Landscape of the protein contact free energy variation upon condensate maturation (300 ns of ageing, $T/T_{c,pure} \sim 0.97$, and bulk condensate density) for the different FUS sequences. $\Delta G/k_BT$ has been obtained from the residue contact probability ratio between aged condensates and liquid-like condensates before ageing. Colour map projections of the free energy landscape in 2-dimensions are also included.

LCD sequences (Figs. 2(c) and 2(d) of the main text) and of the distinct FUS sequences before and after maturation both in presence and absence of RNA (Figure 3(d) and 4(d) of the main text respectively). The methodology for these calculations is described below.

#### A. Estimation of viscosity through the shear relaxation modulus

From NVT simulations, we have evaluated the viscosity within condensates by integrating the relaxation modulus in time (see Chapter 7 of the book [38]):

$$\eta = \int_0^\infty dt\, G(t) \tag{S12}$$

Since our systems are isotropic, we can calculate the shear relaxation modulus $G(t)$ more accurately by using all the components of the stress tensor ($\sigma_{\alpha\beta}$) as shown in Ref. [39]:

$$\begin{aligned} G(t) = \frac{V}{5k_BT}\left[\langle\sigma_{xy}(0)\sigma_{xy}(t)\rangle + \langle\sigma_{xz}(0)\sigma_{xz}(t)\rangle + \langle\sigma_{yz}(0)\sigma_{yz}(t)\rangle\right] \\ + \frac{V}{30k_BT}\left[\langle N_{xy}(0)N_{xy}(t)\rangle + \langle N_{xz}(0)N_{xz}(t)\rangle + \langle N_{yz}(0)N_{yz}(t)\rangle\right], \end{aligned} \tag{S13}$$

where $N_{\alpha\beta} = \sigma_{\alpha\alpha} - \sigma_{\beta\beta}$ is the first normal stress difference. This correlation can be easily calculated by using the compute ave/correlate/long in the USER-MISC package of LAMMPS [40]. In all cases, the relaxation modulus presents an initial regime that mainly accounts for the intramolecular interactions, followed by a terminal region corresponding to much slower relaxation modes, as those coming from intermolecular interactions and the relaxation of the large scale protein and RNA conformations. Due to the very wide range of timescales involved in these calculations and the noisy nature of the relaxation modulus in the terminal region obtained through simulations, we follow a particular route to evaluate the viscosity. At short timescales, $G(t)$ is smooth and the integral can be calculated by applying numerical integration (*i.e.*, trapezoidal rule). However, at longer timescales $G(t)$ is noisier, and hence, we evaluate the integral across that regime by first fitting $G(t)$ to a series of Maxwell modes ($G_i \exp(-t/\tau_i)$) equidistant in logarithmic time [41], and then by calculating the remaining integral analytically. Our fit to the Maxwell modes is carried out with the help of the open-source RepTate software [42]. Finally, viscosity is obtained by adding the two terms:

$$\eta = \eta(t_0) + \int_{t_0}^\infty dt\, G_M(t), \tag{S14}$$

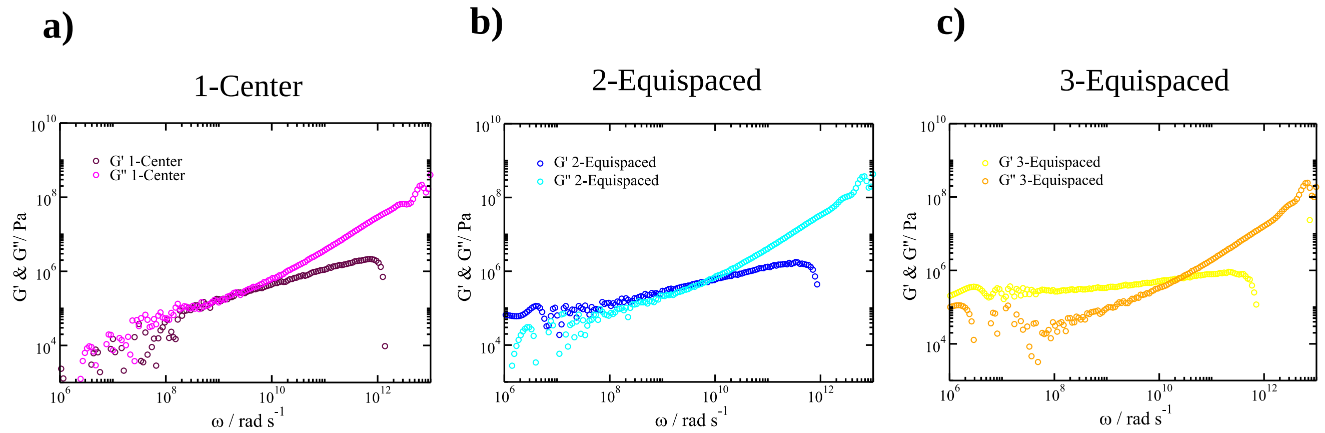


**FIG. S8:** Shear storage modulus (G') and shear loss modulus (G") as a function of frequency ($\omega$) for low-complexity domain phase-separated condensates of the following sequences: 1-Center, 2-Equispaced, and 3-Equispaced. All these calculations were performed after a droplet maturation timescale of 0.1 $\mu$s.

where $\eta(t_0)$ corresponds to the computed term at short time-scales, $G_M(t)$ is the part evaluated via the sum of Maxwell modes fit at long timescales, and $t_0$ is the time that separates both regimes. The error bars present in Fig. 4(d) of the main text have been calculated from the error of the Maxwell mode fits to the value of $G(t)$ obtained in our simulations.

In Fig. S8 we plot G' and G" as a function of $\omega$ to infer whether aged condensates exhibit liquid-like or gel-like behaviour over time. The results of $G'$ and $G''$ from the shear stress relaxation modulus calculations have been obtained by applying the Fourier transform using the open source RepTate software [42]. When G' > G" at short frequencies (i.e., long timescales) elasticity dominates over flow, and gel-like behaviour can be inferred from the plot (i.e., as that shown by the 2-Equispaced and 3-Equispaced sequences). On the other hand, if G" is higher than G', the system exhibits liquid-like behaviour, as shown in the panel of the 1-Center sequence.

### SVIII. SEQUENCE DOMAIN REORDERING OF HNRNPA1

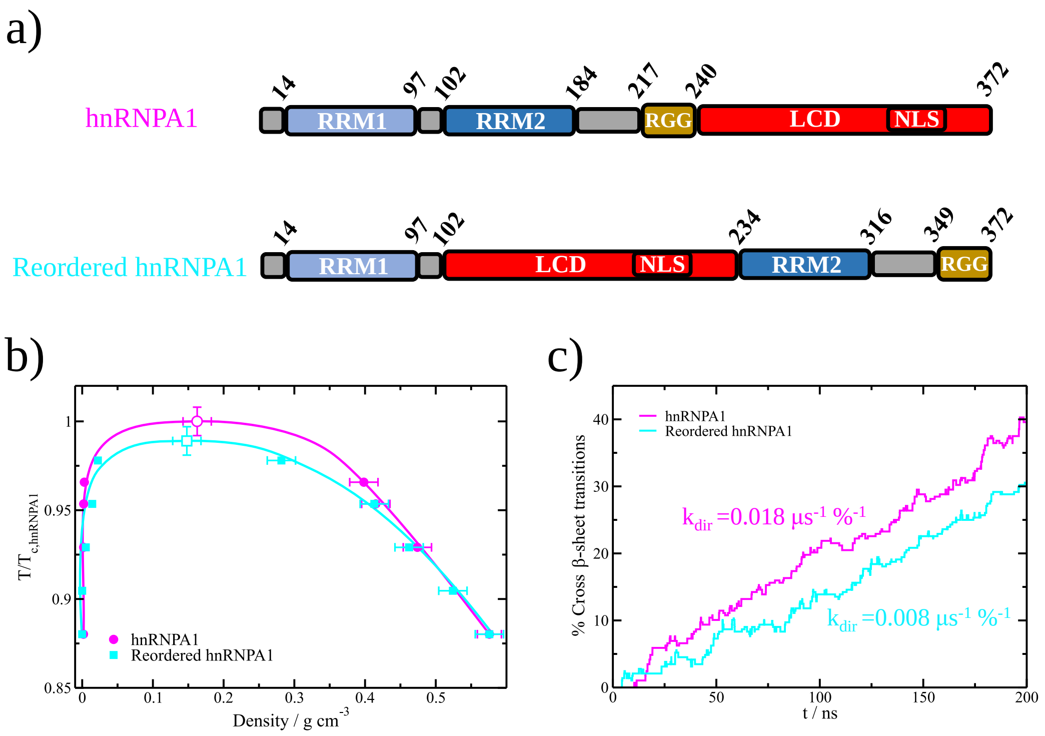


**FIG. S9:** Sequence domain reordering in hnRNPA1 decelerates the accumulation of inter-protein cross-$\beta$-sheets in aged condensates. a) Different domains of the wild-type hnRNPA1 sequence (Top) and the reordered hnRNPA1 variant (Bottom) where LCD accounts for the low-complexity domain, RGG for the Arginine-Glycine-rich region, RRM1 and RRM2 for the RNA-recognition motifs (RRM), and NLS for the nuclear localization sequence. The unique difference between these two sequences is the location of their diverse domains. b) Phase diagram in the $T$-$\rho$ plane for the two studied hnRNPA1 sequences as indicated in the legend. Temperature is renormalized by the critical temperature of pure hnRNPA1 ($T_{c,hnRNPA1}$) which is 409 K. c) Time-evolution of the percentage of LARKS forming inter-peptide cross-$\beta$-sheets in aged condensates of the two hnRNPA1 variants at $T/T_{c,pure}$ ~0.97 (referring $T_{c,pure}$ to the critical $T$ of each hnRNPA1 sequence). The kinetic constant ($k_{dir}$) evaluated through a second-order reaction analysis is provided for each curve.

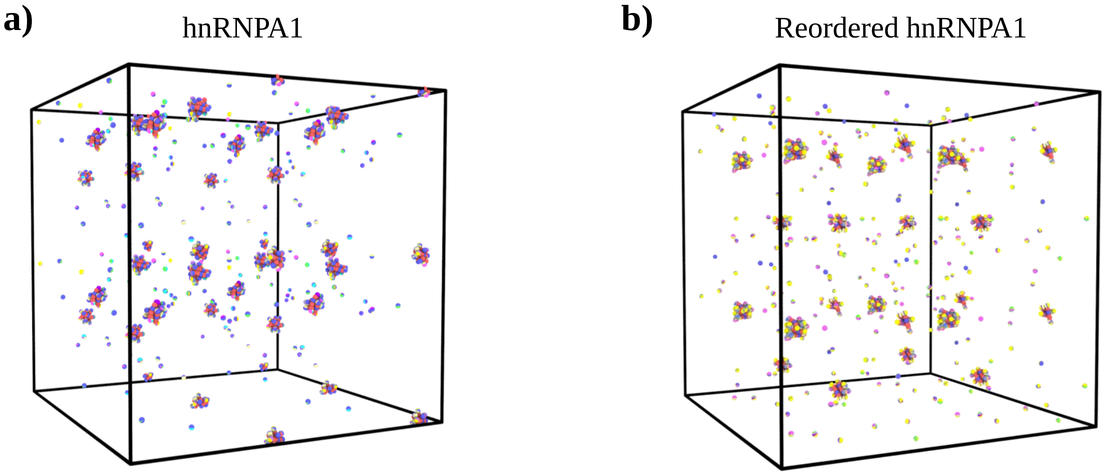


**FIG. S10:** Network connectivity of the different hnRNPA1 sequences calculated with the PPA method after 300 ns of maturation at $T/T_{c,pure}$ ~0.97 within a cubic box at the bulk condensate density at such temperature: a) hnRNPA1 b) Reordered hnRNPA1.

[1] P. Robustelli, S. Piana, and D. E. Shaw, Proceedings of the National Academy of Sciences **115**, E4758 (2018).
[2] H. J. Berendsen, D. van der Spoel, and R. van Drunen, Computer physics communications **91**, 43 (1995).
[3] E. L. Guenther, Q. Cao, H. Trinh, J. Lu, M. R. Sawaya, D. Cascio, D. R. Boyer, J. A. Rodriguez, M. P. Hughes, and D. S. Eisenberg, Nature structural & molecular biology **25**, 463 (2018).
[4] B. Hess, Journal of chemical theory and computation **4**, 116 (2008).
[5] T. Darden, D. York, and L. Pedersen, The Journal of chemical physics **98**, 10089 (1993).
[6] S. Nosé, The Journal of chemical physics **81**, 511 (1984).
[7] M. Parrinello and A. Rahman, Journal of Applied physics **52**, 7182 (1981).
[8] J. S. Hub, B. L. De Groot, and D. Van Der Spoel, Journal of chemical theory and computation **6**, 3713 (2010).
[9] A. Wang, A. E. Conicella, H. B. Schmidt, E. W. Martin, S. N. Rhoads, A. N. Reeb, A. Nourse, D. Ramirez Montero, V. H. Ryan, R. Rohatgi, *et al.*, The EMBO journal **37**, e97452 (2018).
[10] S. Maharana, J. Wang, D. K. Papadopoulos, D. Richter, A. Pozniakovsky, I. Poser, M. Bickle, S. Rizk, J. Guillén-Boixet, T. M. Franzmann, *et al.*, Science **360**, 918 (2018).
[11] A. Garaizar, J. R. Espinosa, J. A. Joseph, and R. Collepardo-Guevara, Scientific reports **12**, 1 (2022).
[12] A. J. Ladd and L. V. Woodcock, Chemical Physics Letters **51**, 155 (1977).
[13] S. Plimpton, Journal of computational physics **117**, 1 (1995).
[14] S. Nosé, The Journal of Chemical Physics **81**, 511 (1984).
[15] W. G. Hoover, Phys. Rev. A **31**, 1695 (1985).
[16] H. Kamberaj, R. Low, and M. Neal, The Journal of chemical physics **122**, 224114 (2005).
[17] S. Das, Y.-H. Lin, R. M. Vernon, J. D. Forman-Kay, and H. S. Chan, Proceedings of the National Academy of Sciences **117**, 28795 (2020).
[18] G. L. Dignon, W. Zheng, Y. C. Kim, R. B. Best, and J. Mittal, PLoS computational biology **14**, e1005941 (2018).
[19] R. M. Regy, G. L. Dignon, W. Zheng, Y. C. Kim, and J. Mittal, Nucleic Acids Research **48**, 12593 (2020).
[20] G. Krainer, T. J. Welsh, J. A. Joseph, J. R. Espinosa, S. Wittmann, E. de Csilléry, A. Sridhar, Z. Toprakcioglu, G. Gudiškytė, M. A. Czekalska, *et al.*, Nature Communications **12**, 1 (2021).
[21] H. S. Ashbaugh and H. W. Hatch, *Journal of the American Chemical Society*, Journal of the American Chemical Society **130**, 9536 (2008).
[22] L. H. Kapcha and P. J. Rossky, Journal of molecular biology **426**, 484 (2014).
[23] S. Plimpton, Journal of Computational Physics **117**, 1 (1995).
[24] T. Schneider and E. Stoll, Physical Review B **17**, 1302 (1978).
[25] R. García Fernández, J. L. F. Abascal, and C. Vega, The Journal of Chemical Physics **124**, 144506 (2006).
[26] J. R. Espinosa, E. Sanz, C. Valeriani, and C. Vega, Journal of Chemical Physics **139** (2013), 10.1063/1.4823499.
[27] J. S. Rowlinson and B. Widom, *Molecular theory of capillarity* (Courier Corporation, 2013).
[28] J. A. Zollweg and G. W. Mulholland, The Journal of Chemical Physics **57**, 1021 (1972).
[29] A. R. Tejedor, A. Garaizar, J. Ramírez, and J. R. Espinosa, Biophysical Journal **120**, 5169 (2021).
[30] A. Garaizar, J. R. Espinosa, J. A. Joseph, G. Krainer, Y. Shen, T. P. Knowles, and R. Collepardo-Guevara, Proceedings of the National Academy of Sciences **119**, e2119800119 (2022).
[31] J. R. Gissinger, B. D. Jensen, and K. E. Wise, Polymer **128**, 211 (2017).
[32] A. R. Tejedor, I. Sanchez-Burgos, M. Estevez-Espinosa, A. Garaizar, R. Collepardo-Guevara, J. Ramirez, and J. R. Espinosa, bioRxiv (2022).
[33] S. V. Kathuria, L. Guo, R. Graceffa, R. Barrea, R. P. Nobrega, C. R. Matthews, T. C. Irving, and O. Bilsel, Biopolymers **95**, 550 (2011).
[34] J. J.-T. Huang, R. W. Larsen, and S. I. Chan, Chemical Communications **48**, 487 (2012).
[35] J. Kubelka, J. Hofrichter, and W. A. Eaton, Current opinion in structural biology **14**, 76 (2004).
[36] A. R. Strom, A. V. Emelyanov, M. Mir, D. V. Fyodorov, X. Darzacq, and G. H. Karpen, Nature **547**, 241 (2017).
[37] S. K. Sukumaran, G. S. Grest, K. Kremer, and R. Everaers, Journal of Polymer Science Part B: Polymer Physics **43**, 917 (2005).
[38] M. Rubinstein, R. H. Colby, *et al.*, *Polymer physics*, Vol. 23 (Oxford university press New York, 2003).
[39] J. Ramírez, S. K. Sukumaran, B. Vorselaars, and A. E. Likhtman, The Journal of chemical physics **133**, 154103 (2010).
[40] S. Plimpton, Journal of computational physics **117**, 1 (1995).
[41] A. E. Likhtman, Macromolecules **38**, 6128 (2005).
[42] V. A. Boudara, D. J. Read, and J. Ramírez, Journal of Rheology **64**, 709 (2020).
